# Vir1p, the Yeast Homolog of Virilizer, is Required for mRNA m^6^A Methylation and Meiosis

**DOI:** 10.1101/2023.02.07.527493

**Authors:** Zachory M. Park, Ethan Belnap, Matthew Remillard, Mark D. Rose

## Abstract

N^6^-Methyladenosine (m^6^A) is one of the most abundant modifications found on eukaryotic mRNAs. mRNA methylation regulates a host of biological processes including meiosis, a specialized diploid cell division program that results in the formation of haploid cells (gametes). During budding yeast meiosis, m^6^A levels peak early, before the initiation of the meiotic divisions. High-throughput studies and work from our lab showed that Ygl036wp, a previously uncharacterized protein interacts with Kar4p, a meiotic protein required for mRNA m^6^A-methylation. Ygl036wp has no discernable domains except for several intrinsically disordered regions. However, protein folding prediction tools showed that Ygl036wp folds like VIRMA/Virilizer/VIR, which is involved in mRNA m^6^A-methylation in higher eukaryotes. In addition, Ygl036wp has several conserved motifs shared with VIRMA/Virilizer/VIR proteins. Accordingly, we propose to call the gene *VIR1* for *budding yeast ortholog of <u>VIR</u>MA/Virilizer/VIR <u>1</u>*. In support, Vir1p interacts with all other members of the yeast methyltransferase complex and is required for mRNA m^6^A methylation and meiosis. Vir1p is required for the stability of proteins comprising the methyltransferase complex, suggesting that Vir1p acts as a scaffold to stabilize the complex. The *vir1*Δ/Δ mutant is defective for premeiotic S-phase, which is suppressed by overexpression of the early meiotic transcription factor *IME1;* additional overexpression of the translational regulator *RIM4* is required for sporulation.

Consistent with *IME1* suppression, *vir1*Δ/Δ exhibits a defect in the abundance of *IME1* mRNA, as well as transcripts within Ime1p’s regulon. Suppression by *IME1* revealed a defect in the expression of the middle meiotic transcription factor, Ndt80p (and genes in its regulon), which is rescued by additional overexpression of *RIM4*. Together, these data suggest that Vir1p is required for cells to initiate the meiotic program and for progression through the meiotic divisions and spore formation.

**Author Summary:** Ygl036wp is a previously uncharacterized protein that we propose to name Vir1p (budding yeast ortholog of <u>VIR</u>MA/Virilizer/VIR <u>1</u>). Work from our lab and others initially found an interaction between Vir1p and members of the yeast mRNA methyltransferase complex (Kar4p and Mum2p). We found that Vir1p interacts with all known members of the methyltransferase complex and is required for mRNA methylation. Vir1p is required early in meiosis; *vir1*Δ/Δ mutants arrest due to the reduced expression of Ime1p. Lower levels of Ime1p cause severe disruption to the meiotic transcriptome in *vir1*Δ/Δ. The *vir1*Δ/Δ meiotic defect can be partially suppressed by the overexpression of *IME1*; full suppression requires overexpression of both *IME1* and *RIM4*. Using recent advances in protein folding predictions, we found that Vir1p is a remote homolog of VIRMA/Virilizer/VIR and shares conserved motifs with the protein from other organisms. Vir1p, like VIRMA/Virilizer/VIR, stabilizes the methyltransferase complex.

## Introduction

mRNA m^6^A methylation is a ubiquitous eukaryotic mRNA modification that has been implicated in a host of important biological processes including spermatogenesis. Mis-regulation of m^6^A methylation has been implicated in a variety of diseases including cancer (Shen, Lan et al. 2020). It is estimated that about 0.1-0.4% of all adenosine nucleotides in mammalian mRNAs are methylated (Yang, Hsu et al. 2018, Zaccara, Ries et al. 2019, Shen, Lan et al. 2020). While the presence of the modification has been known for several decades (Perry and Kelley 1974), questions remain about the underlying mechanisms targeting to specific regions of mRNAs as well as the impact on the downstream fate of the modified mRNAs.

In mammals, mRNA methylation is catalyzed by a core complex that consists of METTL3, METTL14, and WTAP (Bujnicki, Feder et al. 2002, Yang, Hsu et al. 2018, Zaccara, Ries et al. 2019). The core complex is joined by accessory proteins including VIRMA/Virilizer/Vir, HAKAI, RBM15/15B, and ZC3H13. METTL3 is the catalytic subunit and forms a heterodimer with a paralogue, METTL14, which facilitates mRNA binding (Bujnicki, Feder et al. 2002, Liu, Yue et al. 2014, Wang, Doxtader et al. 2016, Wang, Feng et al. 2016). WTAP is thought to link the METTL3/14 heterodimer to accessory proteins (Horiuchi, Kawamura et al. 2013). VIRMA has been implicated in stabilizing the complex and plays a role in site selection of the modification (Yue, Liu et al. 2018). HAKAI is an E3 ubiquitin ligase and its role in the complex is not well understood (Ruzicka, Zhang et al. 2017). RBM15/15B are paralogous RNA-binding proteins that may also be involved in site selection (Lence, Akhtar et al. 2016, Patil, Chen et al. 2016). ZC3H13 links RBM15/15B to WTAP and is responsible for anchoring the complex in the nucleus (Guo, Tang et al. 2018, Knuckles, Lence et al. 2018, Wen, Lv et al. 2018). These proteins make up the “writer” complex, which is responsible for the methylation and the specificity of the modification. Interestingly, METTL3 also engages in a non-catalytic function that involves interactions with Poly-A Binding Protein (PABP) and translation initiation factors to enhance the translation of mRNAs in an m^6^A independent manner (Lin, Choe et al. 2016, Wei, Huo et al. 2022).

In addition to these “writer” proteins, there are several proteins involved in interpreting (“readers”) and erasing (“erasers”) the modification (Dominissini, Moshitch-Moshkovitz et al. 2012, Zheng, Dahl et al. 2013, Li, Zhao et al. 2014, Zhu, Roundtree et al. 2014, Li, Weng et al. 2017). Mammals utilize five reader proteins: YTHDC1-2 and YTHDF1-3. The YTHDF proteins are cytosolic reader proteins that couple translation of methylated mRNAs to their decay (Wang, Lu et al. 2014, Wang, Zhao et al. 2015, Du, Zhao et al. 2016, Shi, Wang et al. 2017). ALKBH5 is the main demethylase that converts m^6^A back to A and is enriched in the nucleus. It is still not clear whether there is active removal of m^6^A outside of the nucleus (Zheng, Dahl et al. 2013).

The mRNA m^6^A methylation complex in the budding yeast *Saccharomyces cerevisiae* is conserved with the higher eukaryotic proteins. The yeast complex contains Ime4p (homolog of METTL3), Kar4p (homolog of METTL14), Mum2p (homolog of WTAP), and Slz1p (homolog of ZC3H13) (Agarwala, Blitzblau et al. 2012). Recent work identified the cytoplasmic dynein light chain, Dyn2p, as being involved in mRNA methylation (Ensinck, Maman et al. 2023 manuscript in preparation). Unlike in higher eukaryotes, mRNA methylation is not found in mitotic mRNA. However, like other eukaryotes, mRNA methylation regulates meiosis in yeast (Clancy, Shambaugh et al. 2002, Agarwala, Blitzblau et al. 2012, Schwartz, Agarwala et al. 2013).

The meiotic program in yeast results in the formation of four quiescent haploid spores and is initiated by starvation for nitrogen and a fermentable carbon source, leading to the expression of the early meiotic transcription factor, *IME1*. Ime1p activates the expression of early genes required for pre-meiotic DNA synthesis and the subsequent initiation of meiotic recombination (Kassir, Granot et al. 1988, Smith and Mitchell 1989, Smith, Su et al. 1990, Smith, Driscoll et al. 1993, Mandel, Robzyk et al. 1994, Kassir, Adir et al. 2003, Neiman 2011). Ime1p also induces the expression of Ime2p, a kinase that has a host of important meiotic functions including facilitating pre-meiotic DNA synthesis and activating the full expression of the middle meiotic transcription factor, Ndt80p (Mitchell, Driscoll et al. 1990, Yoshida, Kawaguchi et al. 1990, Kominami, Sakata et al. 1993, Smith, Driscoll et al. 1993, Foiani, Nadjar-Boger et al. 1996, Dirick, Goetsch et al. 1998, Guttmann-Raviv, Boger-Nadjar et al. 2001, Benjamin, Zhang et al. 2003, Purnapatre, Gray et al. 2005, Schindler and Winter 2006, Sedgwick, Rawluk et al. 2006, Sawarynski, Kaplun et al. 2007, Brush, Najor et al. 2012). Ndt80p regulates the expression of genes required for completion of meiotic recombination and progression through the meiotic divisions as well as spore maturation (Chu, DeRisi et al. 1998, Chu and Herskowitz 1998, Pak and Segall 2002, Kassir, Adir et al. 2003, Shubassi, Luca et al. 2003, Sopko and Stuart 2004, Gurevich and Kassir 2010, Winter 2012). Another key regulator of meiosis is the RNA-binding protein, Rim4p. Rim4p was first identified as an activator of the initial events of meiosis (Soushko and Mitchell 2000, Deng and Saunders 2001), but its role as a negative regulator of translation during meiosis is better understood (Berchowitz, Gajadhar et al. 2013, Berchowitz, Kabachinski et al. 2015, Jin, Zhang et al. 2015). Rim4p forms amyloid-like aggregates that bind transcripts that are produced before their protein products are required (Berchowitz, Kabachinski et al. 2015). Ime2p phosphorylates these aggregates to cause their dissolution, releasing the bound transcripts making them accessible for translation, and allowing the cells to progress through the second meiotic division (Jin, Zhang et al. 2015, Carpenter, Bell et al. 2018, Wang, Zhang et al. 2020).

In yeast, m^6^A levels peak before initiation of the meiotic divisions (Agarwala, Blitzblau et al. 2012). The m^6^A modification is concentrated near stop codons and is associated with translating ribosomes (Bodi, Button et al. 2010, Schwartz, Agarwala et al. 2013, Bodi, Bottley et al. 2015). Similar to mammalian systems, the yeast reader protein Pho92/Mrb1 appears to facilitate both the translation and decay of transcripts important for early meiotic processes, including recombination (Scutenaire, Plassard et al. 2022, Varier, Sideri et al. 2022). One of the best studied examples of the impact of mRNA methylation is the negative transcriptional regulator of *IME1*, *RME1*. Methylation of *RME1* mRNA results in more rapid decay; the subsequent reduction in Rme1p levels facilitates the efficient expression of *IME1* and the induction of the meiotic program (Bushkin, Pincus et al. 2019). However, the methyltransferase complex appears to also have a non-catalytic function that is required for licensing the cells to fully complete meiotic recombination and the divisions (Bushkin, Pincus et al. 2019, Park, Remillard et al. 2023, Park, Sporer et al. 2023). The mechanism of this function in yeast has not yet been fully elucidated but may impact the translation of transcripts in an m^6^A independent manner, similar to the mammalian complex.

Recent work revealed that the yeast karyogamy protein and homolog of METTL14, Kar4p, is required for efficient mRNA methylation and meiosis (Park, Sporer et al. 2023 and Ensinck, Maman et al. 2023 manuscript in preparation). A screen for separation of function alleles of Kar4p identified three distinct functions, one in mating (Mat) (Kurihara, Stewart et al. 1996, Lahav, Gammie et al. 2007) and two in meiosis termed Mei and Spo (Park, Sporer et al. 2023). Loss of Kar4p can be partially suppressed by the overexpression of *IME1* and fully suppressed by the co-overexpression of *IME1* and *RIM4*. *IME1* overexpression permits pre-meiotic DNA synthesis in *kar4*Δ/Δ, but defects remain in the timing of meiotic recombination, the expression of *NDT80*, and proteins required for spore formation. *IME1* and *RIM4* co-overexpression permits sporulation in not only *kar4*Δ/Δ, but also *mum2*Δ/Δ and *ime4*Δ/Δ as well. Interestingly, overexpression of *IME1* alone rescues the meiotic defect of a catalytically dead mutant of Ime4p, as well as *slz1*Δ/Δ (Park, Sporer et al. 2023). Suppression of the Ime4p catalytic mutant by Ime1p overexpression suggests that this “Mei” function is associated with m^6^A methylation. The defects remaining in the deletion mutants after *IME1* overexpression, which required Rim4p overexpression to be suppressed presumably are associated with the non-catalytic function of the complex. Finally, suppression of *slz1*Δ/Δ by *IME1* overexpression alone suggests that the non-catalytic function occurs in the cytoplasm, given Slz1p’s role in anchoring the complex in the nucleus.

The conservation of this complex across eukaryotes led us to hypothesize that there may be other members of the methyltransferase complex in yeast that have yet to be identified. Immunoprecipitation-mass spectrometry (IP-MS) detected a previously uncharacterized protein, Ygl036wp, which interacts with Kar4p and all other known members of the yeast methyltransferase complex. Moreover, Ygl036wp is required for mRNA m^6^A methylation, stability of the complex, and meiosis. Using protein folding prediction algorithms, Ygl036wp is seen to be a remote homolog of VIRMA/Virilizer/VIR. We therefore propose naming this gene *VIR1* for *budding yeast ortholog of <u>VIR</u>MA/Virilizer/VIR <u>1</u>*. Given that many of the mammalian methyltransferase components are essential (Yue, Liu et al. 2018, Poh, Mirza et al. 2022), these findings position yeast as an excellent model for furthering our understanding of mRNA m^6^A methylation.

## Results

### Vir1p Interacts with Kar4p in a Function Specific Manner

Previous work from our lab identified function-specific alleles of Kar4p that are defective either in mating (Mat) or in one or both of its functions in meiosis (Mei and Spo). The Mei function involves Kar4p’s role in mRNA methylation and the Spo function appears to involve a non-catalytic function that promotes the translation of transcripts during meiosis (Park, Sporer et al. 2023, Park, Remillard et al. 2023). To understand the full complement of proteins involved in mRNA methylation and Kar4p’s Spo function, we conducted immunoprecipitation followed by mass spectrometry (IP-MS) to identify meiotic binding partners. We identified two known Kar4p interactors, the components of the methyltransferase complex, Ime4p and Mum2p, as well as a heat shock protein, Ssa4p. Heat shock proteins are common contaminants of this type of experiment. The interaction with Sip5p was not confirmed (data not shown). In addition, we detected a robust interaction with a previously uncharacterized protein called Ygl036wp (Table 1). High-throughput studies had also reported the interaction between Ygl036wp and Kar4p, as well as between Ygl036wp and Mum2p (Ito, Chiba et al. 2001, Gavin, Bosche et al. 2002, Yu, Braun et al. 2008). Using epitope-tagged versions of Ygl036wp and Kar4p, we confirmed the interaction between the two proteins by co-immunoprecipitation (Fig 1A, S Fig 1).

**Table 1.** Top meiotic Kar4p interactors detected by IP-MS.

| Protein | Kar4p Enrichment | Log<sub>2</sub> Normalization | P-value |
| --- | --- | --- | --- |
| Mum2p | 112.5 | 5.2 | 3.92E-05 |
| Kar4p | 85 | 4.8 | 1.24E-04 |
| Ygl036wp/Vir1p | 65.5 | 4.5 | 3.38E-04 |
| Ssa4p | 43.5 | 3.9 | 1.41E-03 |
| Sip5p | 21 | 2.8 | 1.18E-02 |
| Ime4p | 15 | 2.3 | 2.63E-02 |

**Fig 1.**
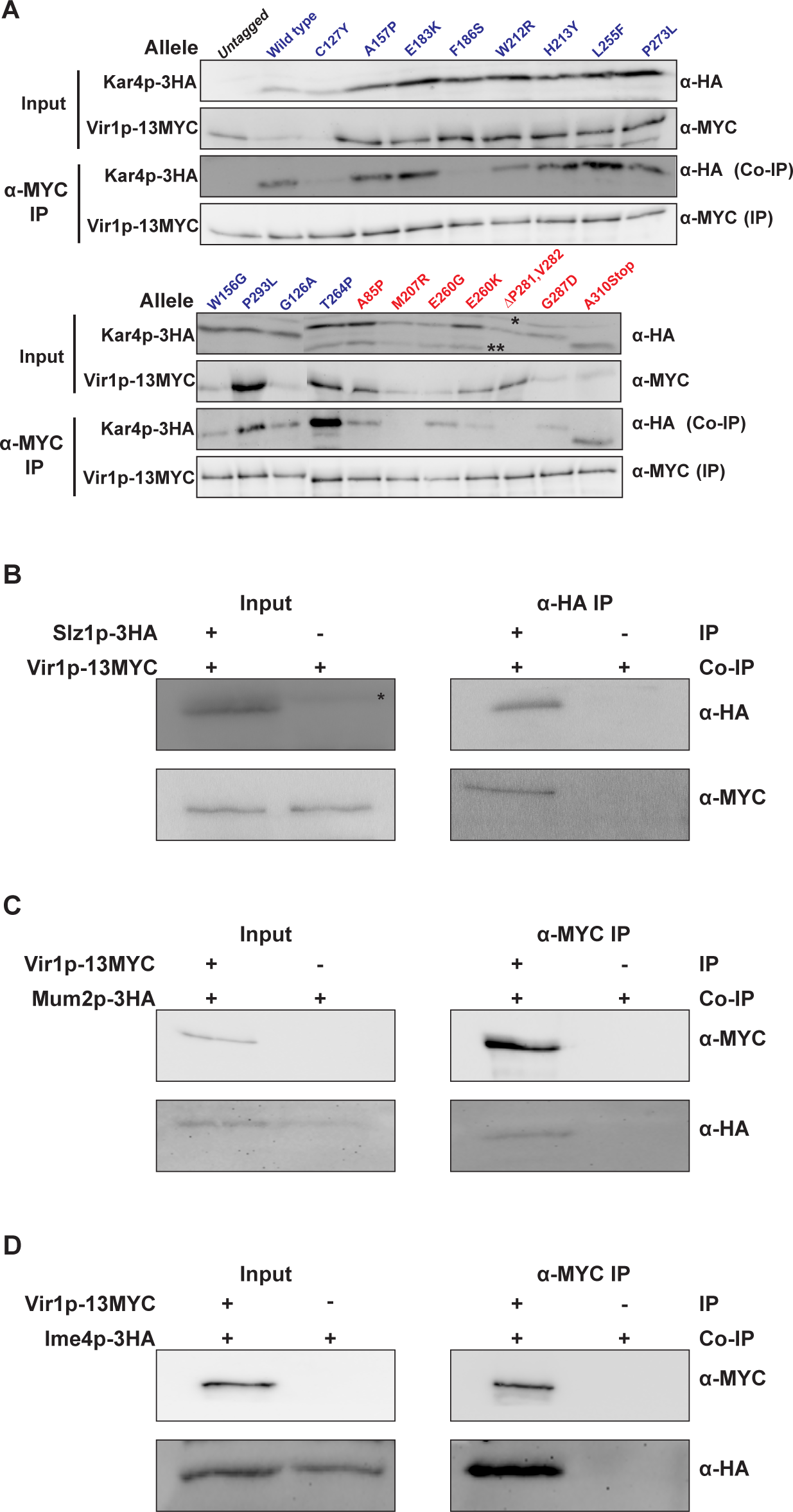
Vir1p engages in a function specific interaction with Kar4p and interacts with all members of the mRNA methyltransferase complex. (A) Western blots of total protein (Input) and Co-IPs (α-MYC IP) between Vir1p-13MYC and mutant alleles of Kar4p-3HA. (Input)Total protein samples from the extracts that were used for the Co-IPs. “*” indicates a non-specific band. “**” indicates degradation products of Kar4p. Note that A310Stop results in the production of a truncated protein. Alleles proficient for Kar4p’s meiotic function (Mei^+^) are in blue and alleles in red are not (Mei^-^). (α-MYC IP) Co-IPs where Vir1p-13MYC was purified and the co-purification of Kar4p-3HA was assayed. Each row is derived from a single blot. One lane with an incorrect sample was cropped from the bottom set of mutation alleles. (B) Western blots of total protein and Co-IPs between Slz1p-3HA and Vir1p-13MYC. (Input) Total protein samples from the extracts that were used for the Co-IPs. “*” indicates a non-specific band. (α-HA IP) Co-IPs where Slz1p-3HA was purified and the co-purification of Vir1p-13MYC was assayed. (C and D) Western blots of total protein and Co-IPs between Mum2p-3HA or Ime4p-3HA and Vir1p-13MYC. (Input) Total protein samples from the extracts that were used for the Co-IPs. (α-MYC IP) Co-IPs where Vir1p-13MYC was purified and the co-purification of either Mum2p-3HA or Ime4p-3HA was assayed.

With the interaction between Ygl036wp and Kar4p validated, we next asked if the two proteins engage in a function-specific interaction, as was seen for other members of the complex (Park, Sporer et al. 2023). Co-immunoprecipitation (Co-IP) was used to assess the interaction between Ygl036wp and the previously identified alleles of Kar4p. Most of the Kar4p alleles that retain meiotic function (blue) maintain the interaction with Ygl036wp, except for C127Y and F186S which have a weak interaction (Fig 1A). Most alleles with defects in meiosis show reduced interaction (red). Remarkably, the pattern of interactions observed between Ygl036wp and the Kar4p mutant proteins matches perfectly the interactions seen with Mum2p, suggesting that Ygl036wp may also play an important role in mRNA methylation. In mammalian systems, the ortholog of Mum2p, WTAP, acts as a linker between the catalytic complex (METTL3/14) and accessory proteins (Horiuchi, Kawamura et al. 2013). These findings suggest that Mum2p may be acting similarly in yeast to bridge the catalytic components (Ime4p and Kar4p) to other components (Ygl036wp and Slz1p). Given the correlation between the interactions of Mum2p and Ygl036wp with the Kar4p alleles and work discussed below, we propose to name Ygl036wp budding yeast ortholog of <u>VIR</u>MA/Virilizer/VIR<u>1</u> (Vir1p).

### Vir1p Interacts with All Members of the mRNA Methyltransferase Complex

Given that Vir1p and Kar4p interact during meiosis, when Kar4p also interacts with other members of the methyltransferase complex, and that high-throughput studies reported an interaction between Vir1p and Mum2p (Gavin, Bosche et al. 2002, Yu, Braun et al. 2008), we determined whether Vir1p interacts with the other members of the methyltransferase complex. By Co-IP, we found that Vir1p binds to all other previously known members of the methyltransferase complex (Slz1p, Mum2p, and Ime4p) (Fig 1B-D).

### Vir1p is Required for mRNA Methylation

Vir1p’s interaction with most of the known mRNA methyltransferase complex members suggested that it might have a role in mRNA methylation. To increase the synchrony and efficiency of meiosis we examined m^6^A in the SK1 strain background (Bodi, Button et al. 2010, Agarwala, Blitzblau et al. 2012, Schwartz, Agarwala et al. 2013, Bodi, Bottley et al. 2015, Bushkin, Pincus et al. 2019), in which we introduced a *vir1*Δ/Δ mutation. Bulk mRNA m^6^A methylation was measured after four hours in sporulation media, which had been previously shown to be when m^6^A levels peaked (Park, Sporer et al. 2023). As expected, m^6^A levels were highly reduced in *vir1*Δ/Δ (Fig 2A), similar to *ime4* Δ/Δ.

**Fig 2.**
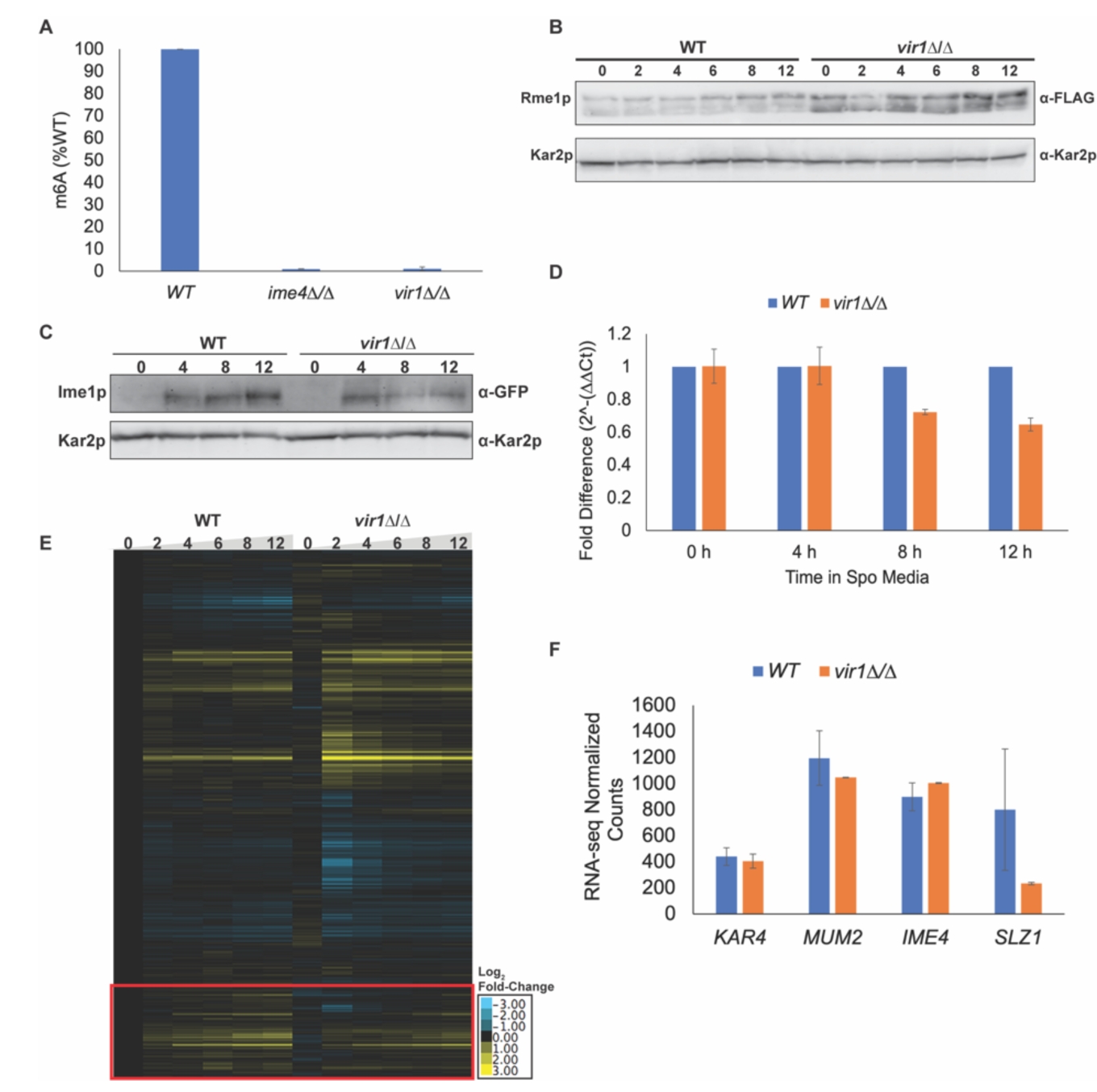
Vir1p is required for mRNA m^6^A methylation. (A) mRNA m^6^A levels measured using an ELISA like assay from EpiGenTek. The indicated mutations were made in the SK1 strain background and samples were harvested after four hours of exposure to meiosis inducing conditions. Experiments were run in three biological replicates for each strain and error bars represent standard deviation. (B) Western Blot of 3xFLAG-Rme1p in wild type and *vir1*Δ/Δ across a time course of meiosis. Kar2p is used as a loading control. (C) Western blot showing GFP-Ime1p levels across a time course of meiosis in wild type and *vir1*Δ/Δ. Kar2p is used as a loading control. (D) qPCR measuring *IME1* transcript levels in wild type and *vir1*Δ/Δ. Data were normalized to levels of *PGK1* expression. Error bars are SEM of three biological replicates. (E) Heatmap of RNA-seq data across a time course of meiosis (0, 2, 4, 6, 8, and 12 hours) in wild type and *vir1*Δ/Δ. Expression was normalized to wild type pre-induction of sporulation (t=0). Genes were clustered in Cluster3.0 and the heatmaps were constructed with Java TreeView. Red box demarcates clusters of genes with delayed and reduced expression in *vir1*Δ/Δ.

Loss of mRNA methylation has been shown to increase levels of the negative regulator of *IME1*, Rme1p. Using an epitope-tagged version of Rme1p, Rme1p levels were increased at least 2-fold in *vir1*Δ/Δ compared to wild type across the time course of meiosis (Fig 2B). Consistent with increased Rme1p levels, there was a ∼2-fold decrease in the level of *IME1* transcript and protein in *vir1*Δ/Δ at 8 and 12 hours in meiosis-inducing conditions (Fig 2C and Fig 2D). This is consistent with work that has shown there is an early burst of *IME1* expression that is m^6^A independent, but that sustained induction of *IME1* requires mRNA methylation (Bushkin, Pincus et al. 2019).

The mis-regulation of *IME1* expression suggested that there should be defects in the meiotic transcriptome of *vir1*Δ/Δ. Using RNA-seq across the time course of meiosis in wild type and *vir1*Δ/Δ, we saw that the transcript profiles were quite similar (Fig 2E). However, a cluster of genes show delayed and reduced expression in the mutant relative to wild type (Fig 2E). To confirm that the impacted genes are involved in meiosis, we did a pairwise comparison to determine differentially expressed genes between wild type and *vir1*Δ/Δ at each time point analyzed. Interestingly, at 2 hours many genes involved in translation and ribosome biogenesis are down regulated in *vir1*Δ/Δ. The reason for this reduction is unclear and warrants further investigation into a potential role for mRNA methylation in the response to glucose starvation alone. Looking specifically at genes with a 2-fold or greater defect at 4 hours in *vir1*Δ/Δ, which is when a gene expression defect becomes noticeable on the heat map, GO-term analysis revealed a specific defect in meiotic progression (Fig 2E red box). Of the 73 genes reduced at 4 hours, the top three GO-terms were “biological process” (p = 0.05), “meiotic cell cycle” (p = 5.2×10^-7^), and “reciprocal meiotic recombination” (p = 4.3×10^-8^). The reduction in early meiotic gene expression is consistent with increased Rme1p and decreased Ime1p levels observed in *vir1*Δ/Δ (Fig 2B and Fig 2C). There was no impact on the expression of other methyltransferase complex members in *vir1*Δ/Δ except for *SLZ1* (Fig 2F). This is expected given that *SLZ1* expression is Ime1p dependent. Together, these data support Vir1p being a member of the mRNA methyltransferase complex and important for the regulation of genes required early in meiosis including *IME1*.

### *vir1*Δ/Δ Mutants Arrest Early in Meiosis

The loss of m^6^A and reduction in Ime1p in *vir1*Δ/Δ suggested that *vir1*Δ/Δ mutants would be unable to sporulate and should arrest early in meiosis before pre-meiotic DNA synthesis. Using wild type and *vir1*Δ/Δ strains expressing *GFP-TUB1* and *SPC42-mCherry*, we used fluorescence microscopy to track meiotic progression (Fig 3A). Wild type strains progressed through both meiotic divisions and reached ∼30% sporulation by 48 hours after induction. In contrast, *vir1*Δ/Δ mutants were arrested with a monopolar spindle and do not progress to sporulation (Fig 3A). To assess the ability of these strains to undergo pre-meiotic DNA synthesis, we used flow cytometry to measure DNA content. Wild type strains began replicating DNA by 12 hours and contained a distinct 4N population by 48 hours. The *vir1*Δ/Δ mutants did not initiate pre-meiotic DNA synthesis and remained arrested at 2N (Fig 3B). This early arrest phenotype is consistent with a role for Vir1p in regulating meiotic entry via mRNA methylation.

**Fig 3.**
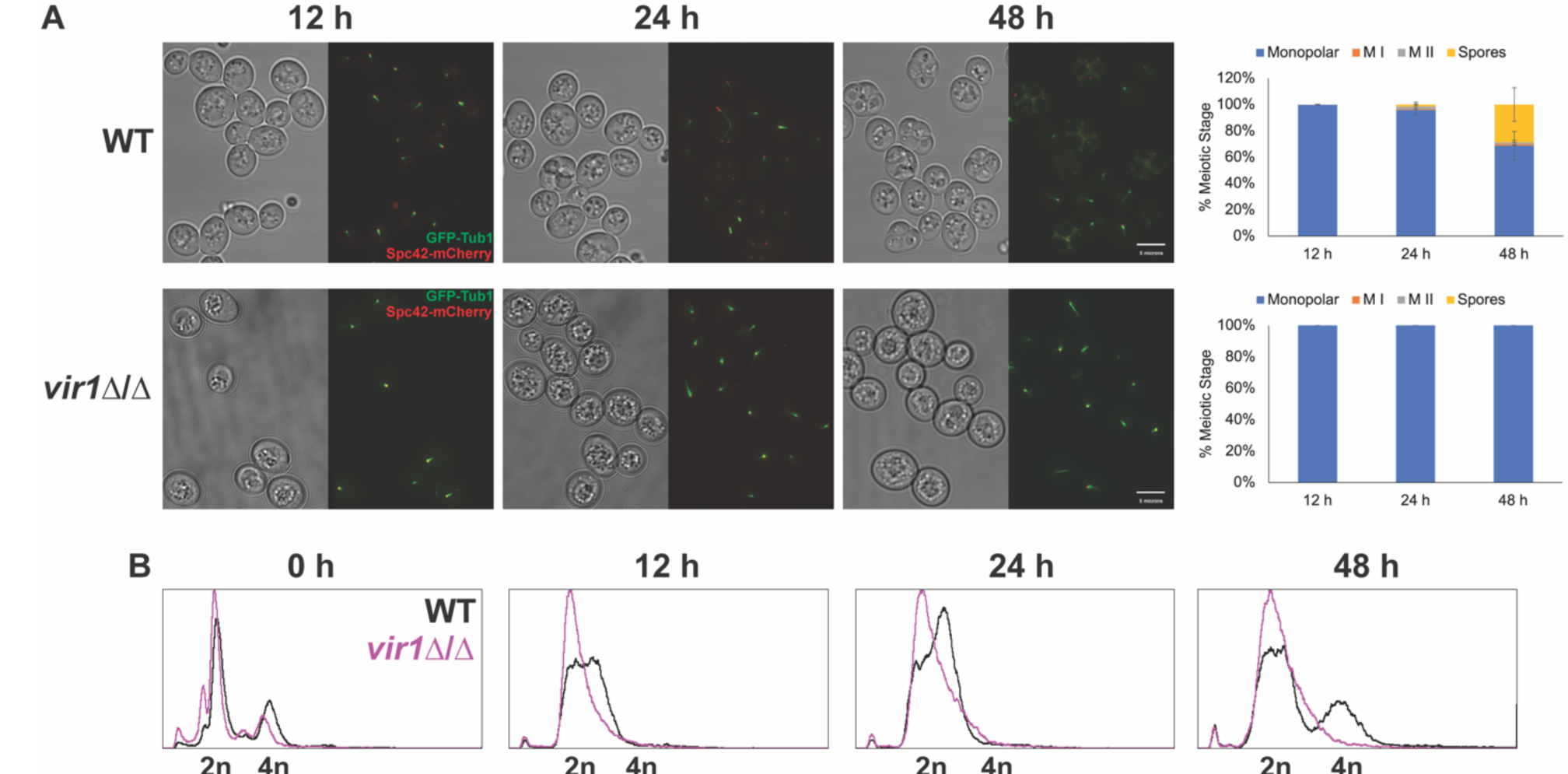
Vir1p is required early in meiosis. (A) Fluorescence microscopy of the spindle pole body (Spc42p-mCherry) and microtubules (GFP-Tub1p) across a time course of meiosis (12, 24, and 48 hours post transfer into sporulation media) in wild type and *vir1*Δ/Δ. Graphs show quantification of the number of cells in different meiotic stages (Monopolar Spindle, Meiosis I, Meiosis II, and Spores). Experiments were run in three biological replicates for each strain and at least 100 cells were counted for each replicate. Error bars represent standard deviation. (B) Flow cytometric analysis of DNA content in wild type and *vir1*Δ/Δ across the same meiotic time course of the microscopy. DNA was stained with propidium iodide.

### Loss of Rme1p and *IME1* Overexpression Partially Suppresses the *vir1*Δ/Δ Meiotic Defect

*RME1* transcripts are one of the key targets of mRNA methylation and methylation leads to their rapid turnover (Bushkin, Pincus et al. 2019). Strains lacking either Kar4p or Ime4p in which Rme1p levels are reduced or absent do progress through pre-meiotic DNA synthesis, but not do not complete meiosis and spore formation (Bushkin, Pincus et al. 2019, Park, Sporer et al. 2023). Suppression of Kar4p’s Mei function by loss of Rme1p was due to the restoration of normal levels of *IME1* expression (Park, Sporer et al. 2023). Given that Vir1p also appears to be working in this same pathway, we asked if a *vir1*Δ/Δ *rme1*Δ/Δ double mutant could undergo pre-meiotic DNA synthesis. By flow cytometry, otherwise wild type strains lacking Rme1p began to form a 4N population faster than in strains with Rme1p. As expected for mutations in the methyltransferase complex, *vir1*Δ/Δ *rme1*Δ/Δ double mutants underwent pre-meiotic DNA synthesis. However, these strains still failed to sporulate (Fig 4A and S Fig 2).

**Fig 4.**
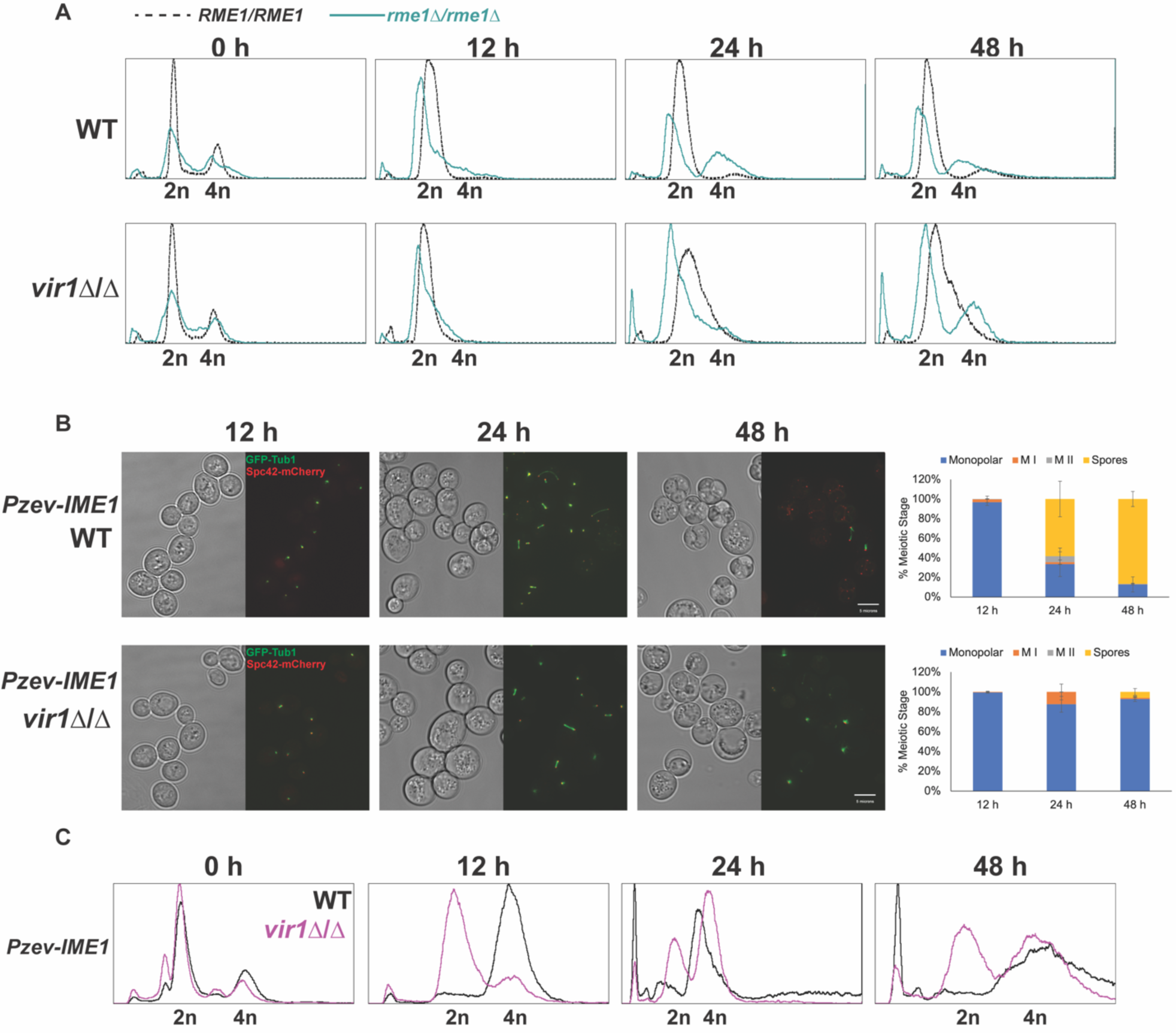
Loss of Rme1p and overexpression of *IME1* partially rescues the *vir1*Δ/Δ meiotic defect. (A) Flow cytometric analysis of DNA content in wild type and *vir1*Δ/Δ carrying the wild type allele of *RME1* or a deletion of *RME1* across a time course of meiosis. DNA was stained with propidium iodide. (B) Fluorescence microscopy of the spindle pole body (Spc42p-mCherry) and microtubules (GFP-Tub1p) across a time course of meiosis (12, 24, and 48 hours post movement into sporulation media) in wild type and *vir1*Δ/Δ with *IME1* overexpressed from an estradiol-inducible promoter. Graphs are the quantification of the number of cells in different meiotic stages (Monopolar Spindle, Meiosis I, Meiosis II, and Spores). Experiments were run in three biological replicates for each strain and at least 100 cells were counted for each replicate. 1 µM of estradiol was used to induce expression. Error bars represent standard deviation. (C) Flow cytometric analysis of DNA content in wild type and *vir1*Δ/Δ with *IME1* overexpressed across the same meiotic time course of the microscopy. DNA was stained with propidium iodide.

Given that loss of the negative regulator of *IME1*, Rme1p, partially suppressed the *vir1*Δ/Δ meiotic defect, we asked directly if overexpression of *IME1* also suppressed, as it does with *kar4*Δ/Δ (Park, Sporer et al. 2023, Park, Remillard et al. 2023). *IME1* was overexpressed using the estradiol inducible P_Z3EV_ promoter system (Pzev) (McIsaac, Gibney et al. 2014). Wild type strains sporulated very efficiently after *IME1* overexpression, reaching 86% sporulation by 48 hours. However, 93% of the *vir1*Δ/Δ mutants were arrested with a monopolar spindle; only 6% formed spores after *IME1* overexpression (Fig 4B). As in *kar4*Δ/Δ, *IME1* overexpression permitted pre-meiotic DNA synthesis in *vir1*Δ/Δ, more robustly than the *vir1*Δ/Δ *rme1*Δ/Δ double mutant (Fig 4C). This suggests that loss of Rme1p can facilitate some level of meiotic entry in *vir1*Δ/Δ, but either the level of Ime1p is insufficient or there are still other factors inhibiting the mutant. The requirement for these other factors is only fully bypassed upon the direct overexpression of *IME1*.

To understand the lack of sporulation after *IME1* overexpression in *vir1*Δ/Δ, we used RNA-seq to look at the transcriptome. The overexpression of *IME1* revealed later transcript level defects in both the expression of the middle meiotic transcription factor, *NDT80*, and Ndt80p-dependent genes (Fig 5A (red box) and Fig 5B). Some 58% of the genes highlighted by the red box in Figure 5A were found to be regulated by Ndt80p. This finding is consistent with data for *kar4*Δ/Δ after *IME1* overexpression, indicating that the block in meiosis after *IME1* overexpression in *vir1*Δ/Δ is also upstream of *NDT80* expression (Park, Remillard et al. 2023). Loss of Rme1p (Bushkin, Pincus et al. 2019) and *IME1* overexpression alone (Park, Sporer et al. 2023) can rescue the meiotic defect of a catalytically dead mutant of Ime4p, suggesting that remaining defects after overexpression of *IME1* do not involve mRNA methylation. Thus, Vir1p may also be involved in the potential non-catalytic function of the complex.

**Fig 5.**
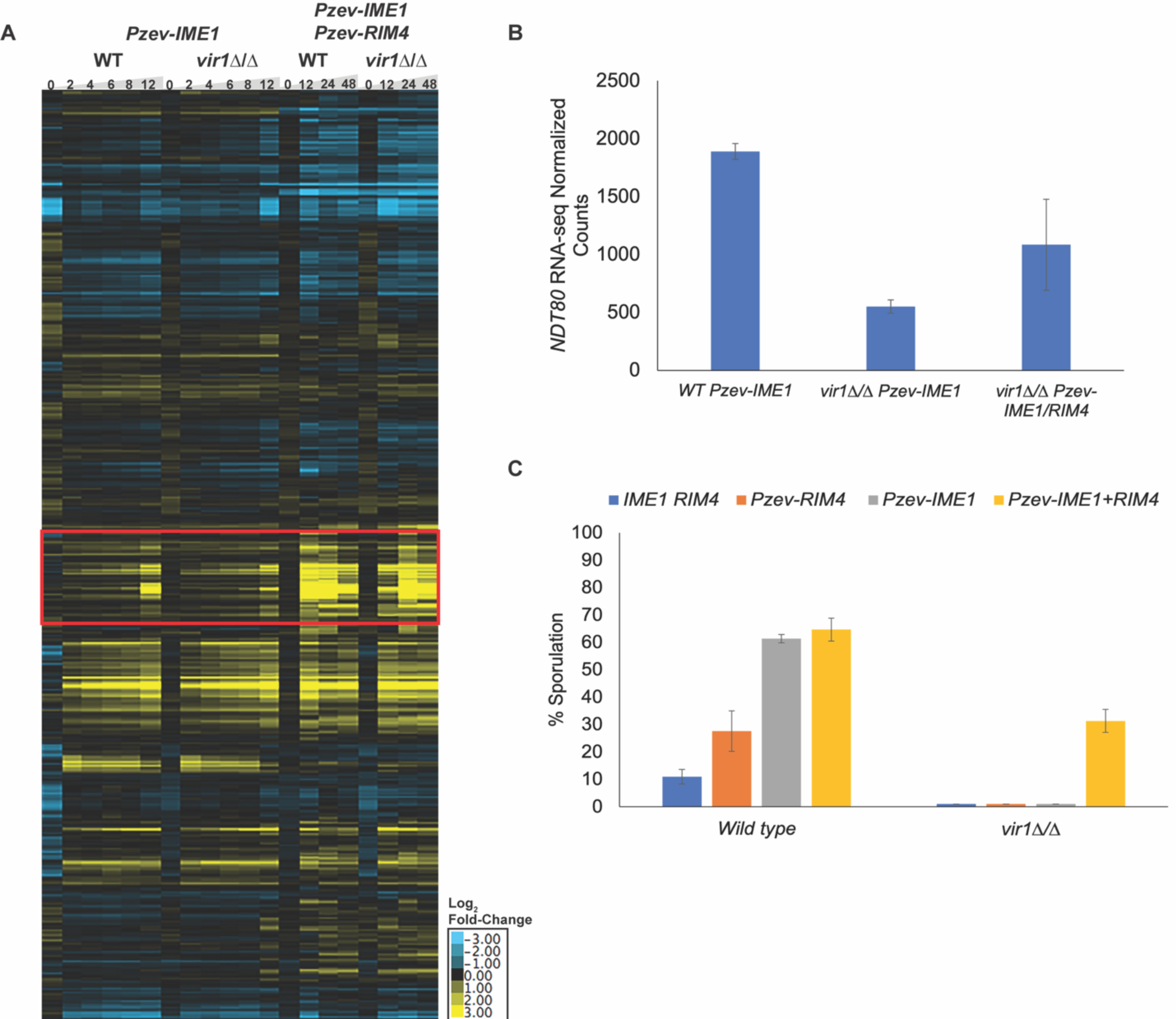
Co-overexpression of *IME1* and *RIM4* rescues the *vir1*Δ/Δ meiotic defect. (A) Heatmap of RNA-seq data across a time course of meiosis in wild type and *vir1*Δ/Δ with *IME1* overexpressed and *IME1*/*RIM4* co-overexpressed. Expression was normalized to wild type t=0 used in Fig 2. Genes were clustered in Cluster3.0 and the heatmaps were constructed with Java TreeView. Red box demarcates cluster of genes with delayed and reduced expression in *vir1*Δ/Δ. (B) *NDT80* RNA-seq normalized counts from wild type and *kar4*Δ/Δ with *IME1* overexpressed and from *kar4*Δ/Δ with *IME1* and *RIM4* overexpressed. Counts were normalized using the standard normalization method in DESeq2. Error bars represent standard deviation between two biological replicates. (C) Sporulation of wild type and *vir1*Δ/Δ with either endogenous expression of *IME1* and *RIM4*, overexpression of *RIM4*, overexpression of *IME1*, or overexpression of both *IME1* and *RIM4*. 1 µM of estradiol was used to induce expression. All dyads, triads, and tetrads were counted across three biological replicates for each strain. At least 100 cells were counted after 48 hours post addition of estradiol. Error bars represent standard error of three biological replicates.

### *IME1* and *RIM4* Overexpression Suppresses the *vir1*Δ/Δ Meiotic Defect

The co-overexpression of *IME1* and the translational regulator, *RIM4*, fully suppresses the meiotic defect of *kar4*Δ/Δ, *mum2*Δ/Δ, and *ime4*Δ/Δ (Park, Sporer et al 2023, Park, Remillard et al. 2023). Given the similarities between these mutants and *vir1*Δ/Δ, we tested whether *IME1* and *RIM4* co-overexpression also fully rescues the *vir1*Δ/Δ meiotic defect. Using strains with both *Pzev-IME1* and *Pzev-RIM4*, wild type strains sporulate more efficiently with either or both genes overexpressed (Fig 5C). The *vir1*Δ/Δ mutant also sporulated after co-overexpression, but not as efficiently, and delayed by 24-hours compared to wild type, suggesting that defects persist in this strain (Fig 5C and S Fig 3). However, co-overexpression did restore *vir1*Δ/Δ to roughly the same level and timing of sporulation as wild type strains with *RIM4* overexpressed (Fig 5C and S Fig 3). To understand why there is still a delay in sporulation after *IME1* and *RIM4* co-overexpression, we again used RNA-seq. Even with co-overexpression, there was a pronounced delay in *NDT80* expression in *vir1*Δ/Δ, as was seen in *kar4*Δ/Δ under the same conditions (Fig 5B) (Park, Remillard et al. 2023). The persistent delay in *NDT80* expression explains the delayed and inefficient sporulation in *vir1*Δ/Δ.

### Vir1p is Required for Efficient Expression of Multiple Meiotic Proteins

The requirement for *RIM4* overexpression for the completion of meiosis suggested that *vir1*Δ/Δ mutants, like *kar4*Δ/Δ mutants, would have defects in the levels of key meiotic proteins (Park, Remillard et al. 2023). The level of the meiotic kinase Ime2p was not reduced in *kar4*Δ/Δ with endogenous *IME1* expression, indicating that Ime2p is not limiting the progression of *kar4*Δ/Δ mutants through the early stages of meiosis (Park, Remillard et al. 2023). However, after overexpression of *IME1* in *kar4*Δ/Δ, the level of Ime2p was found to be reduced later in meiosis, compared to wild type (Park, Remillard et al. 2023). In *vir1*Δ/Δ mutants without *IME1* overexpression, *IME2* transcript levels were reduced about 2-fold, but Ime2p levels were normal (S Fig 4). The reduction in *IME2* transcript is consistent with the reduction in Ime1p expression, although the lack of an effect on the level of the protein is surprising. After *IME1* overexpression, there was a ∼2-fold reduction in *IME2* transcript and a similar reduction in Ime2p levels in *vir1*Δ/Δ (Fig 6A and Fig 6B), similar to *kar4*Δ/Δ. Co-overexpression of *IME1* and *RIM4* in *vir1*Δ/Δ both increased Ime2p levels and reduced the time to peak expression compared to *IME1* overexpression alone (Fig 6C). It is likely that the delay in *NDT80* expression in *vir1*Δ/Δ after *IME1* overexpression (Fig 5B) leads to the loss of the late induction of *IME2* expression, as previously reported in *NDT80* mutants (Shin, Skokotas et al. 2010).

**Fig 6.**
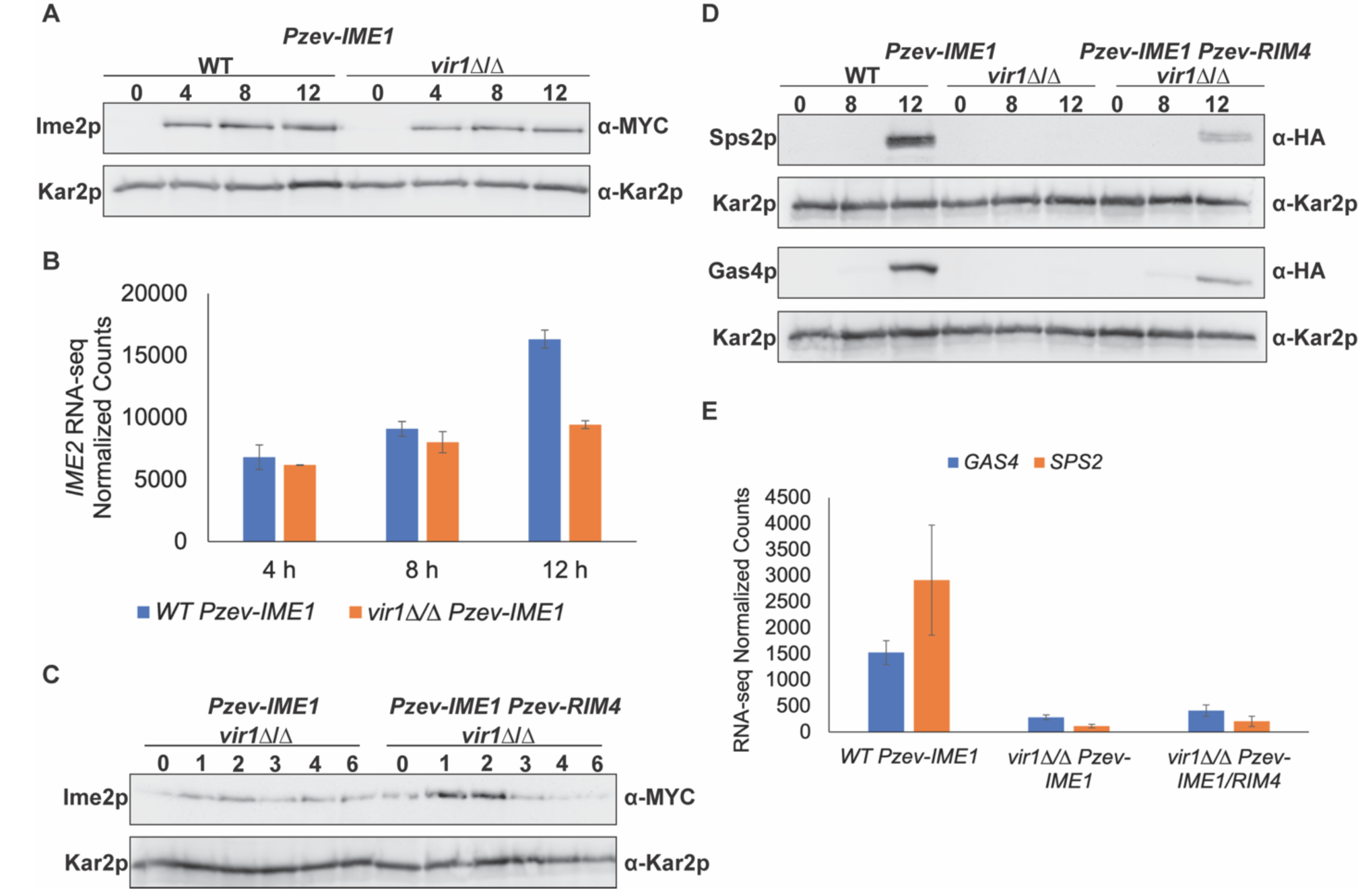
Vir1p is required for the full expression of Ime2p, Sps2p, and Gas4p. (A) Western blots of Ime2p-13MYC across a meiotic time course in wild type and *vir1*Δ/Δ with *IME1* overexpressed. Kar2p is used as a loading control. (B) *IME2* RNA-seq counts normalized between wild type and *kar4*Δ/Δ with *IME1* overexpressed. Counts were normalized using the standard normalization method in DESeq2. Error bars represent standard deviation between two biological replicates. (C) Western blots of Ime2p-13MYC across a time course of meiosis in *vir1*Δ/Δ with either *IME1* overexpressed or *IME1* and *RIM4* overexpressed. Kar2p is used as a loading control. (D) Western blots of Sps2p-3HA and Gas4p-3HA in wild type and *vir1*Δ/Δ with *IME1* overexpressed and in *vir1*Δ/Δ with *IME1* and *RIM4* overexpressed. Blots in the same row were exposed for the same length of time. Kar2p serves as a loading control. (E) *SPS2* (Top) and *GAS4* (Bottom) RNA-seq normalized counts from 12-hour time points in wild type with *IME1* overexpressed and *vir1*Δ/Δ with either *IME1* overexpressed or *IME1* and *RIM4* overexpressed. Counts were normalized using the standard normalization method in DESeq2. Error bars represent standard deviation between two biological replicates.

Mass spectrometry of the meiotic proteome with *IME1* overexpressed revealed other proteins with low levels in *kar4*Δ/Δ, including the mid-late meiotic proteins Gas4p and Sps2p (Park, Remillard et al. 2023). Similarly, in *vir1*Δ/Δ, *GAS4* and *SPS2* transcript and protein levels were very low after *IME1* overexpression compared to wild type (Fig 6D and Fig 6E). Overexpression of *IME1* and *RIM4* only slightly increased transcript levels but caused a large increase in protein levels. The persistent delay in *NDT80* expression explains the low transcript and protein levels of *GAS4* and *SPS2,* given that they are Ndt80p-dependent genes (Fig 5B, Fig 6D and Fig 6E). Nevertheless, overexpression of Rim4p greatly increased the levels of both Gas4p and Sps2p. Rim4p was first identified as an early activator of meiotic gene expression, which may be related to the increased levels of Gas4p and Sps2p levels. Thus, in *vir1*Δ/Δ, as in *kar4*Δ/Δ, defects in the expression of proteins upstream of *NDT80* effect the expression of proteins needed later in the meiotic program (Park, Remillard et al. 2023), which are not rescued by overexpressing *IME1*. Taken together, these data show that Vir1p behaves like other methyltransferase complex members by affecting regulation both upstream and downstream of Ime1p.

### Vir1p is the Yeast Homolog of VIRMA/Virilizer/VIR

Given the high degree of conservation between the core components the of mRNA methyltransferase complex in eukaryotes, we asked if there are also homologs of Vir1p. Vir1p is highly conserved amongst the ascomycetes, however, no obvious homologs were found in outgroups. Vir1p has several predicted disordered regions suggesting that the selection on maintaining primary amino acid sequence is not as strong (S Fig 5A). However, remote homologs have been identified that share structure and function, but are not strongly conserved at the primary amino acid sequence (Holm 2022). The yeast homologs of many components of the methyltransferase complex in other eukaryotes including METTL3 (Ime4p), METTL14 (Kar4p), WTAP (Mum2p), and Z3CH13 (Slz1p) are already known. There are only three other members of the complex that have been identified in eukaryotes thus far: HAKAI, RMB15/15B, and VIRMA/Virilizer/VIR. The Alpha Fold predictions of Vir1p, were compared to the predicted structures of the three proteins that have yet been shown to have a yeast homolog. Of the three, the predicted structure of Vir1p was strongly similar to that of VIRMA/Virilizer/VIR.

To quantify the similarities in the predicted structures, we used the DALI program (Holm 2022) to determine if Vir1p is predicted to fold like VIRMA. Although VIRMA and related proteins are much larger than Vir1p, and Vir1p lacks an N-terminal beta-pleated sheet domain, there is a core region of all that shares similar predicted structure, whether between Vir1p and human VIRMA (Fig 7A), plants (VIR) (Fig 7B), or flies (Virilizer) (Fig 7C). The similarity to Virilizer was the highest, with a Z-score of 13.9 and an RMSD of 10.1. Comparing Virilizer and VIR to VIRMA resulted in comparable statistics; Virilizer had a Z-score of 24.5 and RMSD of 5.4, while VIR had a Z-score of 14.4 and RMSD of 14.4. The VIRMA-like proteins from all three organisms shared about 10% identity to Vir1p. Similarly, the RaptorX DeepAlign program (Kallberg, Wang et al. 2012) identified the same region of conserved structure between the VIRMA homologs and Vir1p (S Fig 6). Again, Virilizer had the strongest match to Vir1p with an RMSD of 5.63 and a TM-score of 0.426.

**Fig 7.**
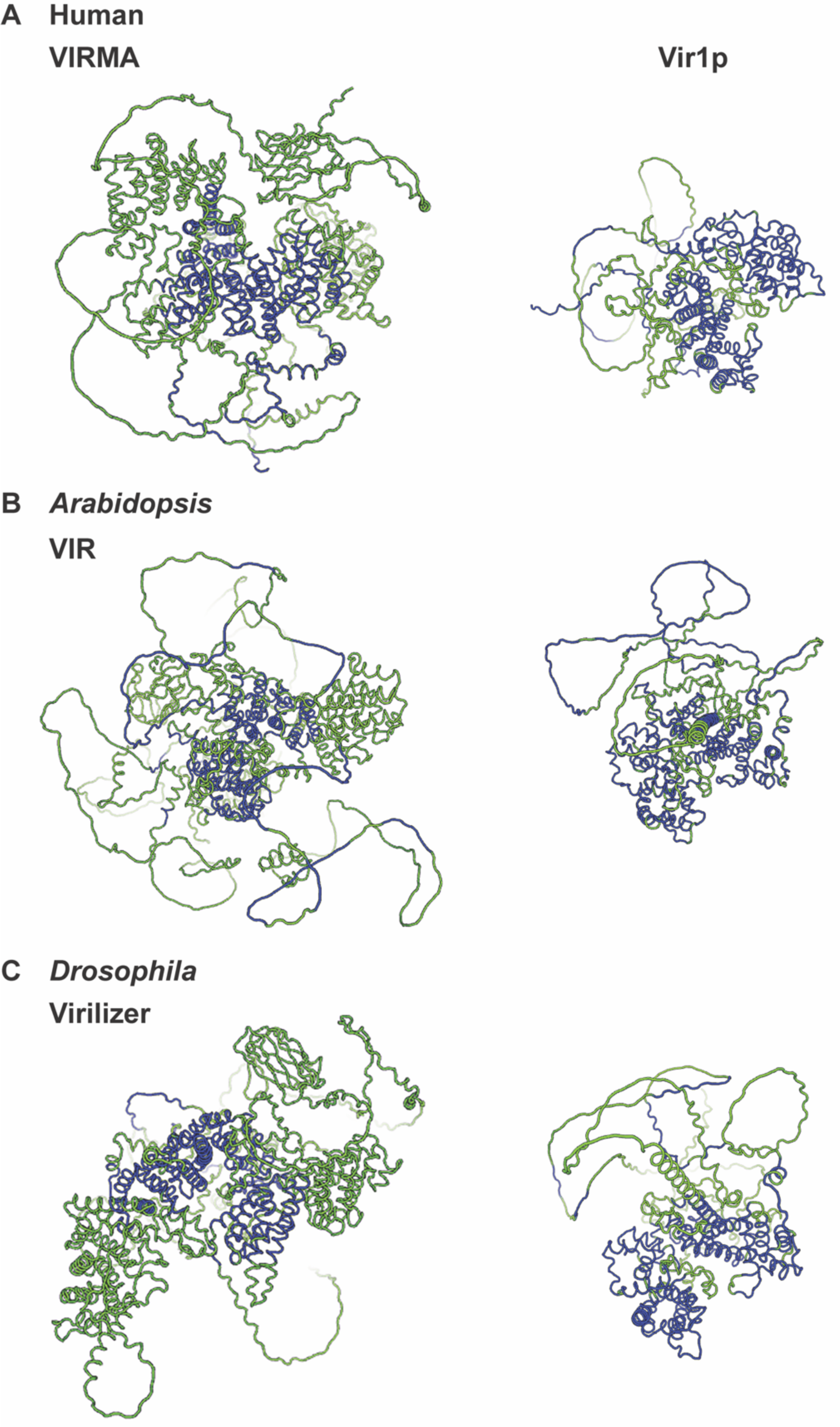
Vir1p is the yeast homolog of VIRMA/Virilizer/VIR. Dali predictions of conserved structure between Vir1p and VIRMA from (A) humans, (B) *Arabidopsis*, and (C) *Drosophila*. (Left) Image of the AlphaFold predicted structure of VIRMA (green) with the region conserved with Vir1p highlighted in blue. (Right) Image of the AlphaFold predicted structure of Vir1p (green) with the region conserved with VIRMA highlighted in blue. The Z-score for each comparison was well above the Z-score of 2, which would be considered spurious similarity.

Interestingly, Blastp revealed conserved motifs located in the disordered C-terminal region of Vir1p (S Fig 5B). DEPICTER (Barik, Katuwawala et al. 2020), which computationally predicts the functions of intrinsically disordered domains, strongly predicts that the C-terminal disordered region of Vir1p is involved in DNA binding and protein-protein interaction, and less strongly in RNA binding. It is tempting to speculate that Vir1p’s disordered C-terminus may function to anchor the complex to chromatin while it methylates mRNA. Published data suggests that other complex members do not associate with chromatin (Ensinck, Maman et al. 2023 manuscript in preparation). Thus, it is more likely that Vir1p’s disordered regions are important for facilitating protein-protein interactions.

### Vir1p is Required for the Stability of the mRNA Methyltransferase Complex

A second requirement for identifying proteins with remote homology is that they share similar functions. Research on VIRMA suggests that it acts to stabilize other complex members including WTAP (Yue, Liu et al. 2018). If Vir1p is a remote homolog of VIRMA, then Vir1p should also be required for the stability of the complex. The stability of Kar4p, Mum2p, and Ime4p was measured during meiosis in *vir1*Δ/Δ using cycloheximide to block new protein synthesis. All three proteins were unstable in *vir1*Δ/Δ. The turnover rate of Kar4p was 2-fold faster in *vir1*Δ/Δ compared to wild type (Fig 8A). Mum2p is very stable in wild type with a half-life longer than 60 minutes, but in *vir1*Δ/Δ the half-life dropped to ∼24 minutes (Fig 8B). Ime4p was so unstable in *vir1*Δ/Δ that it could not be detected, preventing measurement of the half-life in the mutant (Fig 8C). We did not examine the stability of Slz1p because we saw strong defects in *SLZ1* transcript levels in *vir1*Δ/Δ (Fig 2F).

**Fig 8.**
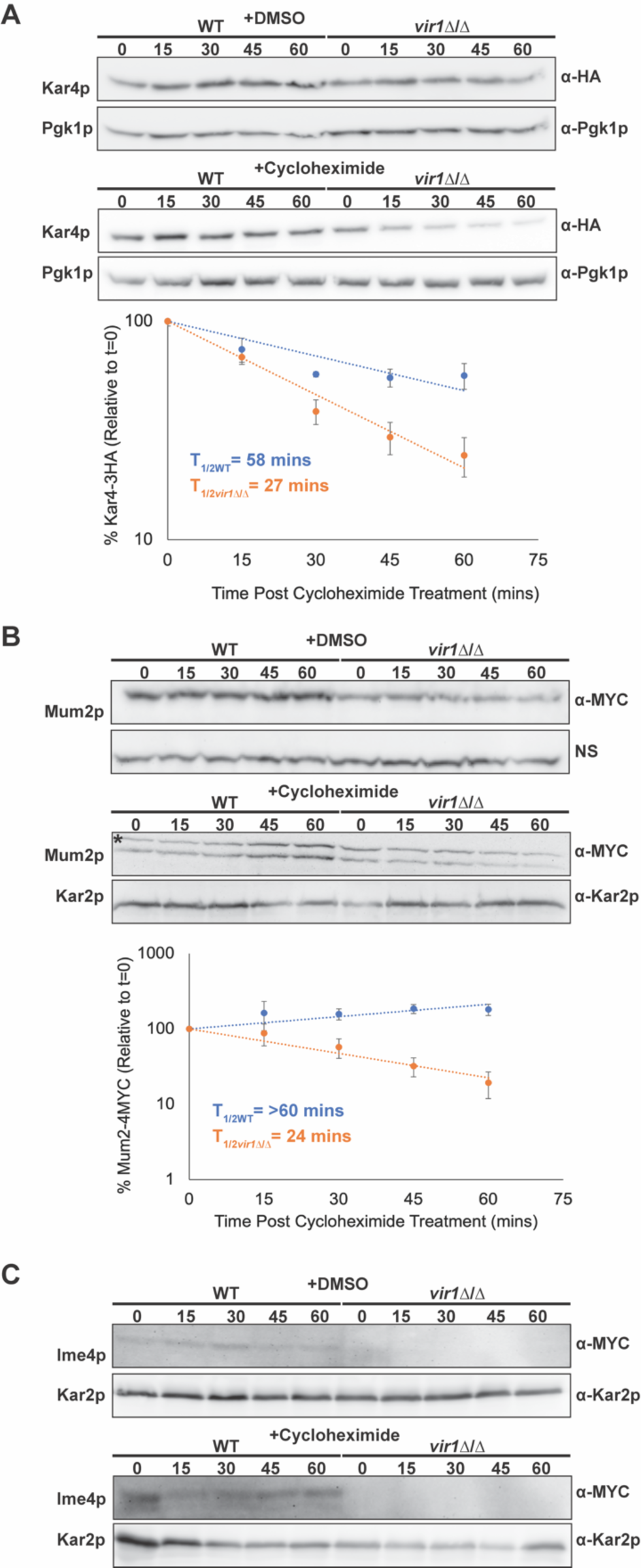
Vir1p is required for the stability of the mRNA methyltransferase complex. (A) Western blots of Kar4p-3HA after four hours in meiosis inducing media with either 100 µM cycloheximide or an equivalent amount of DMSO in both wild type and *vir1*Δ/Δ. (Top) Kar4p-3HA levels with DMSO. (Bottom) Kar4p-3HA levels with cycloheximide. Graph is the quantification of three biological replicates of the cycloheximide chase experiment. Pgk1p is used as a loading control. (B) Western blot of Mum2p-4MYC (bottom band) after four hours in meiosis inducing media with either 100 µM cycloheximide or an equivalent amount of DMSO in both wild type and *vir1*Δ/Δ. (Top) Mum2p-4MYC levels with DMSO. A non-specific band is used as a loading control (Bottom) Mum2p-4MYC levels with cycloheximide. Graph is the quantification of three biological replicates of the cycloheximide chase experiment. Kar2p is used as a loading control. “*” indicates a non-specific band. Error bars in A and B are SEM of the three biological replicates. (C) Western blot of 3MYC-Ime4 after four hours in meiosis inducing media with either 100 µM cycloheximide or an equivalent amount of DMSO in both wild type and *vir1*Δ/Δ. (Top) 3MYC-Ime4 levels with DMSO. (Bottom) 3MYC-Ime4 levels with cycloheximide. Kar2p is used as a loading control.

We next examined the impact of deleting other complex members on the stability of the overall complex. We previously found that Ime4p levels are not reduced in *kar4*Δ/Δ (Park, Sporer et al. 2023). Vir1p and Mum2p levels were also not impacted in *kar4*Δ/Δ (S Fig 7A and S Fig 7B). However, the absence of Ime4p resulted in significantly lower levels of Kar4p, and Kar4p turned over faster in *ime4*Δ/Δ (S Fig 8A). This is further evidence that Kar4p’s role in meiosis is dependent on its interaction with Ime4p. Mum2p levels were also reduced in *ime4*Δ/Δ, and the turnover rate was slightly increased, although not to the same extent as for Kar4p (S Fig 8B). Neither the levels of Vir1p nor its turnover were impacted in *ime4*Δ/Δ (S Fig 8C). A concurrent paper (Ensinck, Maman et al. 2023, manuscript in preparation) found that Vir1p levels were lower in *ime4*Δ/Δ in the SK1 strain background. The discrepancy is most likely due to differences in meiotic efficiency between SK1 and the S288c strain background used in this study. SK1 undergoes meiosis much more rapidly than S288c, and so the defect in Vir1p levels may arise at later points in meiosis. In *mum2*Δ/Δ, similar to the effect of *vir1*Δ/Δ, the turnover rate of Kar4p was increased though not to the same extent as in *vir1*Δ/Δ and Ime4p was not detectable (S Fig 9A, S Fig 9C). The overall level of Vir1p was slightly reduced in *mum2*Δ/Δ, but there was no impact on Vir1p’s turnover rate (S Fig 9B).

**Fig 9.**
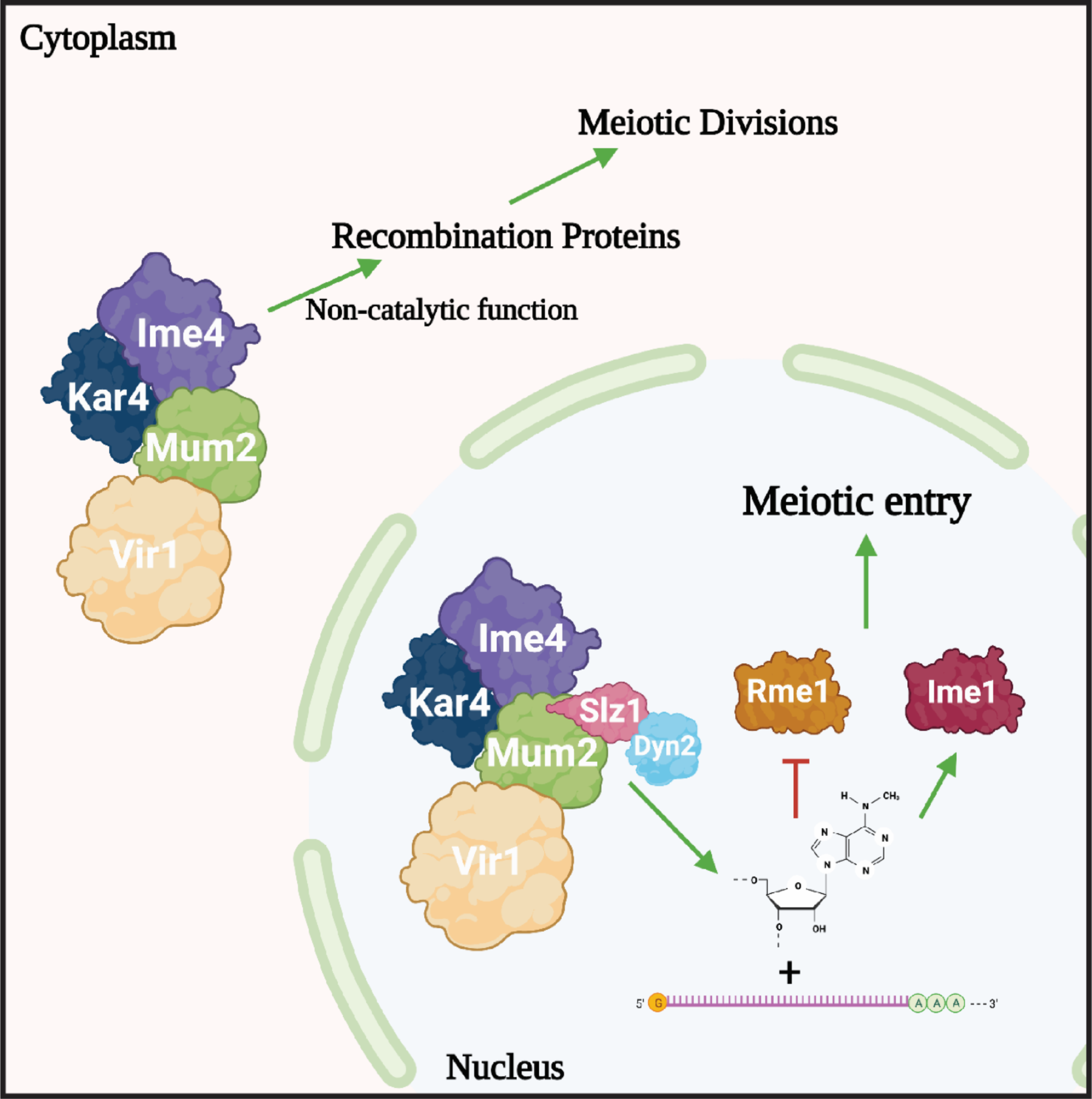
A model for the function of Vir1p in meiosis. Vir1p functions in mRNA m^6^A methylation with Ime4p, Mum2p, Kar4p, Slz1p and Dyn2p. mRNA methylation facilitates meiotic entry through regulation of Rme1p and Ime1p. Vir1p, Kar4p, Mum2p, and Ime4p engage in a function seemingly independent of m^6^A catalysis that is important for the progression into the meiotic divisions. Figure was created with BioRender.com.

Taken together, while removal of some complex members results in reduced levels of other members, only removal of Vir1p increased the turnover rate of all other complex members, consistent with a scaffolding role stabilizing the methyltransferase complex. Thus, the conservation of both structure and function with VIRMA/Virilizer/VIR support Vir1p being a remote homolog of those proteins.

## Discussion

The initial characterization of the mRNA m^6^A methyltransferase complex in yeast suggested that it comprised only Ime4p, Mum2p, and Slz1p (Agarwala, Blitzblau et al. 2012). Subsequent work indicated that proteins found in the complex in other eukaryotes are conserved in yeast, such as Kar4p. Recent work from our lab (Park, Sporer et al. 2023) and others (Ensinck, Maman et al. 2023 manuscript in preparation) showed that the complex also contains Kar4p. Work presented here and by Ensinck et al. (2023 manuscript in preparation) show that Vir1p, the VIRMA/Virilizer/VIR homolog, is also part of the complex. Thus, the yeast methyltransferase complex is much more highly conserved than previously thought.

Building on our findings of the role of Kar4p in meiosis, we found that Vir1p is required early in meiosis and *vir1*Δ/Δ mutants have a severe loss of mRNA m^6^A methylation. The impact on the expression of *IME1* and Ime1p-dependent genes in *vir1*Δ/Δ provides further evidence that mRNA methylation acts upstream of *IME1* to promote meiotic entry. That *IME1* overexpression and loss of Rme1p repression only partially suppress the *vir1*Δ/Δ defect indicate that Vir1p is also involved in other functions of the complex. The role of Vir1p in other functions is characterized by defects in protein levels in *vir1*Δ/Δ which can be bypassed by *RIM4* overexpression. It is probable that, like Kar4p (Park, Remillard et al. 2023), Vir1p is required for the efficient expression of proteins involved earlier in the meiotic program, including in meiotic recombination (Fig 9). The persistent block in meiosis after *IME1* overexpression is likely due to these defects. It will be interesting to determine what exactly remains impacted in these mutants after *IME1* overexpression and whether the yeast complex utilizes a similar mechanism to enhance translation as has been shown in mammalian cells (Lin, Choe et al. 2016, Wei, Huo et al. 2022).

Utilizing the previously identified separation of function mutants of Kar4p (Park, Sporer et al. 2023), we found that mutants that impact the interaction between Mum2p and Kar4p also impact the interaction between Vir1p and Kar4p. This suggests that in yeast, as in mammals (Horiuchi, Kawamura et al. 2013), Mum2p (WTAP) links the catalytic components of the complex Ime4p (METTL3) and Kar4p (METTL14) to regulatory proteins. It will be interesting to determine if the interaction between Slz1p and Dyn2p also matches this pattern. One Kar4p allele, T264P, continues to be interesting given that it is strongly defective for all Kar4p’s meiotic functions, but still interacts with Vir1p, Mum2p, and Ime4p (Park, Sporer et al. 2023). One possibility is that this allele impacts the ability of Kar4p to bind mRNA and future work will investigate that possibility. Identifying an allele that results in a fully formed complex that can’t interact with mRNA could be a valuable tool for determining the underlying mechanism of the additional functions of the complex.

Work on VIRMA in mammalian cells has suggested that it plays a role in concentrating m^6^A around stop codons and in the 3’ UTR (Yue, Liu et al. 2018). However, the mechanism by which it helps confer this specificity is not well understood. The predicted functions for the disordered C-terminal domain makes Vir1p an interesting candidate to determine how the methyltransferase specifically methylates some mRNAs. The disordered region may play a role in engaging the mRNA and facilitating site selection, and so mutations in Vir1p’s C-terminus should alter the distribution of m^6^A. Alternatively, Vir1p’s C-terminal domain might facilitate interaction with a protein or proteins that help confer specificity. In the mammalian complex, there are two paralogous RNA binding proteins, RBM15/15B, that have also been implicated in site selection. Ensinck et al. (2023 manuscript in preparation) identified another component of the complex in yeast, Dyn2p, that is linked to the complex through Slz1p but did not find that it was orthologous to RBM15/15B or any other identified member of the complex in other organisms. There may be other components of the yeast methyltransferase complex that are yet to be identified; characterization of Vir1p’s interactors might identify novel members.

Vir1p’s ability to stabilize the complex might be mediated via the disordered domains acting to facilitate phase separation to sequester the other complex members away from the protein degradation machinery. In the absence of Vir1p, the proteins would not be protected and so rapidly degraded. Work has also shown that methylated mRNAs bound by reader proteins are more likely to undergo phase separation mediated by the low-complexity domains of the reader proteins (Yoon, Ringeling et al. 2017). Regardless of the potential role of Vir1p’s disordered region, Vir1p provides several new avenues to explore mRNA methylation in yeast.

In summary, Vir1p is a conserved component of the yeast methyltransferase complex and is required for at least two steps in meiosis: an early step that is upstream of *IME1* in part through regulation of *RME1* and a later step that is at least partially upstream of *NDT80*. The high degree of conservation between the yeast methyltransferase complex and that of mammals and other eukaryotes positions yeast as an excellent model for furthering our understanding of this important mRNA modification.

## Materials and Methods

### Sporulation

Cells were first grown overnight at 30 °C in either YPD (yeast nitrogen base (1% w/v), peptone (2% w/v), and 2% glucose) or synthetic media lacking a nutrient to maintain selection of a plasmid. Overnight cultures were diluted to an OD of 0.1 in YP-Acetate (yeast nitrogen base (1% w/v), peptone (2% w/v), and potassium acetate (1% w/v)) and grown for 16 to 18 hours at 30 °C. YPA cultures were then moved into 1% (w/v) potassium acetate (Spo) media supplemented with uracil, histidine, and leucine at an OD of 0.5. Cultures were sporulated for various amounts of time at 26 °C. For experiments involving overexpression, estradiol was added to cultures immediately after moving them to Spo media at a final concentration of 1 µM. Experiments using the SK1 strain background were handled in much the same way with the only two exceptions being sporulation cultures were diluted to a final OD of 2.0 and were sporulated at 30 °C.

### Immunoprecipitation-Mass Spectrometry

For each time point, 200 ml of sporulating culture at an OD_600_ of 1 were pelleted, washed with water, and flash frozen by liquid nitrogen. Samples were prepared according to the Thermo Scientific Pierce Magnetic HA-Tag IP/Co-IP Kit (88838). Cells were lysed using a FastPrep (MP) bead beater (5 x 90 second runs at 4°C) and acid-washed glass beads (BioSpec), centrifuged, and lysate prepared according to kit protocol.

Eluted immune-precipitated samples were dried in a speedvac to completion and the residual proteins were resuspended in 200 μl of 50 mM ammonium bicarbonate pH 8. TCEP (tris(2-carboxyethyl)phosphine) was added to a final concentration of 5 mM and incubated at 60°C for 20 min. 15 mM chloroacetamide was added and incubated in the dark at room temperature for a further 30 min. One microgram of Trypsin Gold (Promega) was added to each sample and incubated in an end-over-end mixer at 37°C for 16 hours. An additional 0.25 μg of Trypsin Gold was added and incubated in an end-over-end mixer at 37°C for another 3 hours. Samples were acidified by adding TFA (trifluoroacetic acid) to a final concentration of 0.2% and were desalted using SDB-RPS (styrenedivinyl benzene reverse phase sulfonate) stage-tips (Rappsilber, Mann et al. 2007). Samples were dried completely in a speedvac and resuspended with 20 μl of 0.1% formic acid pH 3. Five microliters were injected per run using an Easy-nLC 1000 UPLC system. Samples were loaded directly onto a 45 cm long 75 μm inner diameter nano capillary column packed with 1.9 μm C18-AQ (Dr. Maisch, Germany) mated to metal emitter in-line with an Orbitrap Elite (Thermo Scientific, USA). The mass spectrometer was operated in data dependent mode with the 120,000 resolution MS1 scan (400-1800 m/z) in the Orbitrap followed by up to 20 MS/MS scans with CID fragmentation in the ion trap. Dynamic exclusion list was invoked to exclude previously sequenced peptides for 120 s if sequenced within the last 30s and maximum cycle time of 3 s was used.

Raw files were searched using MS-Amanda (Bern, Kil et al. 2012) and Sequest HT algorithms (Eng, McCormack et al. 1994) within the Proteome Discoverer 2.1 suite (Thermo Scientific, USA). 15 ppm MS1 and 0.5 Da MS2 mass tolerances were specified. Carbamidomethylation of cysteine was used as a fixed modification, oxidation of methionine, and deamidation of asparagine were specified as dynamic modifications. Trypsin digestion with maximum of 2 missed cleavages were allowed. Files were searched against the yeast SGD database downloaded 13 Jan 2015 and supplemented with common contaminants.

Scaffold (version Scaffold_4.7.5, Proteome Software Inc., Portland, OR) was used to validate MS/MS based peptide and protein identifications. Peptide identifications were accepted if they could be established at greater than 95.0% probability by the Scaffold Local FDR algorithm. Protein identifications were accepted if they could be established at greater than 99.9% probability and contained at least 2 identified peptides. Protein probabilities were assigned by the Protein Prophet algorithm (Nesvizhskii, Keller et al. 2003). Proteins that contained similar peptides and could not be differentiated based on MS/MS analysis alone were grouped to satisfy the principles of parsimony.

### Co-Immunoprecipitation

Cells were sporulated for four hours as described above and a volume of cells equivalent to 25 OD units was harvested for each strain. Cells were resuspended in 200 µL of lysis buffer (1% Triton-X 100, 0.2 M Tris-HCl pH 7.4, 0.3 M NaCl, 20% glycerol, 0.002 M EDTA) supplemented with 10x protease inhibitor (Pierce) and lysed using the FastPrep (MP) for four rounds of 40 seconds at 4.0 m/s with one minute on ice between each round of bead beating.

Protein extracts were then incubated with 15 µL of anti-MYC or HA magnetic beads (Pierce) for one hour at room temperature. Beads were washed three times with lysis buffer, resuspended in deionized water, and moved to a new tube before being resuspended in 2x sample buffer and boiled for 5 minutes to elute proteins off the beads. Samples were then resolved on 8% SDS-PAGE gels and western blotting was conducted as described below.

### Western Blotting

Protein samples were resolved on an appropriate percentage SDS-PAGE gel before being transferred to a PVDF membrane using a semi-dry transfer apparatus (TransBlot SD BioRad) at 25 volts for 30 minutes. After transfer, membranes were blocked for 30 minutes using 10% milk in TBS before being incubated with primary antibody (anti-HA (12CA5) 1:1,000, anti-HA (rabbit: Cell Signaling C29F4) 1:2,500, anti-MYC (9E10) 1:1000, anti-MYC (rabbit: Novus NB600-336SS) 1:1,000, anti-FLAG (Sigma M2) 1:1000, anti-GFP (BD Living Colors 632377) 1:1000, anti-PGK1 (Invitrogen 22C5D8) 1:1000) for one hour at room temperature. Membranes were then washed three times for ten minutes each with 0.1% TBS-Tween20 (TBST) before being incubated with the appropriate secondary antibody in 1% milk in TBST for 30 minutes at room temperature. Incubation with secondary antibody (donkey anti-mouse IgG (Jackson ImmunoResearch) 1:10,000 or donkey anti-rabbit IgG (ImmunoResearch) 1:20,000) was then followed by one brief rinse in TBST and another three washes for ten minutes each with TBST before being incubated with Immobilon Western HRP substrate (Millipore) for 5 minutes and imaged using the G-Box from SynGene. Densitometry was conducted using ImageJ.

### Protein Extraction

Proteins were extracted as previously described (Park, Sporer, et al. 2023). Samples were incubated with 150 µl of 1.85 M NaOH on ice for 10 minutes. 150 µl of 50% TCA was then added to the samples and they were left to incubate at 4°C for 10 minutes. Samples were then washed with acetone before being resuspended in 100 µl of 2x sample buffer. Samples were then boiled for five minutes and then kept on ice for five minutes before being spun down and moved to new tubes. Samples were then either run on an SDS-PAGE gel or stored for later use at -80°C.

### Microscopy

Microscopy was conducted as previously described (Park, Sporer, et al. 2023). Briefly, 100 µl of a sporulated culture expressing GFP-*TUB1* integrated at *TUB1* (Straight, Marshall et al. 1997) and *SPC42*-mCherry was spun down and resuspended in 10 µl of Spo media. Cells were imaged on a DeltaVision deconvolution microscope (Applied Precision, Issaquah, WA) based on a Nikon TE200 (Melville, NY). All images were deconvolved using Applied Precision SoftWoRx imaging software. At least one hundred cells were counted in all cases, and the percentage of cells in each meiotic stage including mature spores (dyads, triads, and tetrads) were determined using the morphology of the spindle and number of spindle pole bodies.

### Flow Cytometry

Flow cytometry was conducted as previously described (Park, Sporer, et al. 2023). Cultures were induced to sporulate and at each time point measured 500 µl of cells were pelleted and washed in dH_2_O. Cells were fixed in 70% ethanol and stored at -20 °C for one hour up to several days. After fixation, cells were washed in 500 µl 50mM sodium citrate (pH 7.2) and incubated in 0.5 ml sodium citrate containing 0.25 mg/ml RNaseA for two hours at 37°C. After RNaseA treatment, 500 µl of 8 µg/mL propidium iodide in sodium citrate buffer was added to each sample and incubated overnight in the dark at 4 °C. The cells were then sonicated at 50% duty cycle, output setting 3, for 4×12 pulses on ice. FACS was conducted at the Flow Cytometry and Cell Sorting Shared Resource core at Georgetown University and data were analyzed using FCS Express.

### mRNA m^6^A Methylation

mRNA methylation levels were measured as previously described (Park, Sporer, et al. 2023). SK1 cultures were sporulated for four hours. Cells were lysed using the FastPrep (MP) as described above. Total RNA was then harvested using the Qiagen RNeasy kit with on-column DNase treatment. mRNA was purified from these total RNA samples using Oligod(T) magnetic beads (Thermofisher). mRNA methylation was measured using the EpiQuik fluorometric m^6^A RNA methylation quantification kit (EpiGenTek). The kit protocol was followed as written. Briefly, purified mRNA was bound to wells and then washed before the addition of an antibody to m^6^A. Wells were then washed again before the addition of a detection antibody followed by an enhancer/developer solution. The fluorescence was measured using a Synergy plate reader (BioTek) at 530_EX_/590_EM_ nm and m^6^A levels were quantified for each sample using a standard curve generated from a positive control provided with the kit. Measurements from an *ime4*Δ/Δ strain were used to subtract background from the data collected from the *vir1*Δ/Δ mutant.

### RNA-seq

RNA-seq was conducted as previously described (Park, Remillard, et al. 2023). Cells were lysed and RNA was purified as described above. RNA samples were sent to Novogene Corporation for library prep and mRNA sequencing using an Illumina based platform (PE150). Resulting data were analyzed using the open access Galaxy platform (Afgan, Baker et al. 2018). Reads were first mapped to the yeast genome (sacCer3) using the standard settings in BWA-MEM and then counted using htseq-count. Differential expression analysis was conducted using DESeq2 and heat maps were made using Cluster 3.0 and Java Tree View. GO-term analysis was conducted using <u>Yeastract</u> (Teixeira, Monteiro et al. 2014).

### qPCR

qPCR was conducted as previously described (Park, Sporer, et al. 2023). RNA was harvested as described above. cDNA libraries were constructed using the High-Capacity cDNA Reverse Transcription kit (Applied Biosystems) with 10 µl of the total RNA sample. The concentration of the resulting cDNA was measured using a nanodrop. qPCR reactions were set up using Power SYBR Green PCR Master Mix (Applied Biosystems) with 50 ng of total RNA. The reactions were run on a CFX96 Real-Time System (BioRad) with reaction settings exactly as described in the master mix instructions with the only change being the addition of a melt curve at the end of the program. Results were analyzed using CFX Maestro. Primer sequences were as follows: *PGK1* Forward 5’-CTCACTCTTCTATGGTCGCTTTC-3’, *PGK1* Reverse 5’-AATGGTCTGGTTGGGTTCTC-3’, *IME1* Forward 5’-ATGGCAACTGGTCCTGAAAG-3’, and *IME1* Reverse 5’-GGAACGTAGATGCGGATTCAT-3’.

### Protein Fold Similarity Predictions

Alpha Fold structure prediction files for Vir1p (AF-P53185) and VIRMA from humans (AF-Q69YN4), *Arabadopsis* (AF-F4J8G7), and *Drosophila* (AF-Q9W1R5) were downloaded from UniProt. The downloaded pdb files were uploaded to the <u>DALI</u> or <u>RaptorX DeepAlign</u> web-based platform to conduct pairwise comparisons between Vir1p and the different versions of VIRMA. Standard input settings were used for all comparisons and the whole protein sequence was used for the predictions.

## Data Availability

Source data for the RNA-seq experiments can be found using GEO ascension number <u>GSE222684</u>.

## Acknowledgements

We would like to thank Anne Rosenwald for helpful feedback on this manuscript. We thank Tharan Srikumar for his expert Mass Spectrometry and Emily Schmidt for technical assistance. We thank Folkert van Werven for providing the plasmid for tagging *IME1* and for sharing information prior to publication. This work was supported by NIH grants GM037739 and GM126998 to MDR.

---

**S Fig 1.**
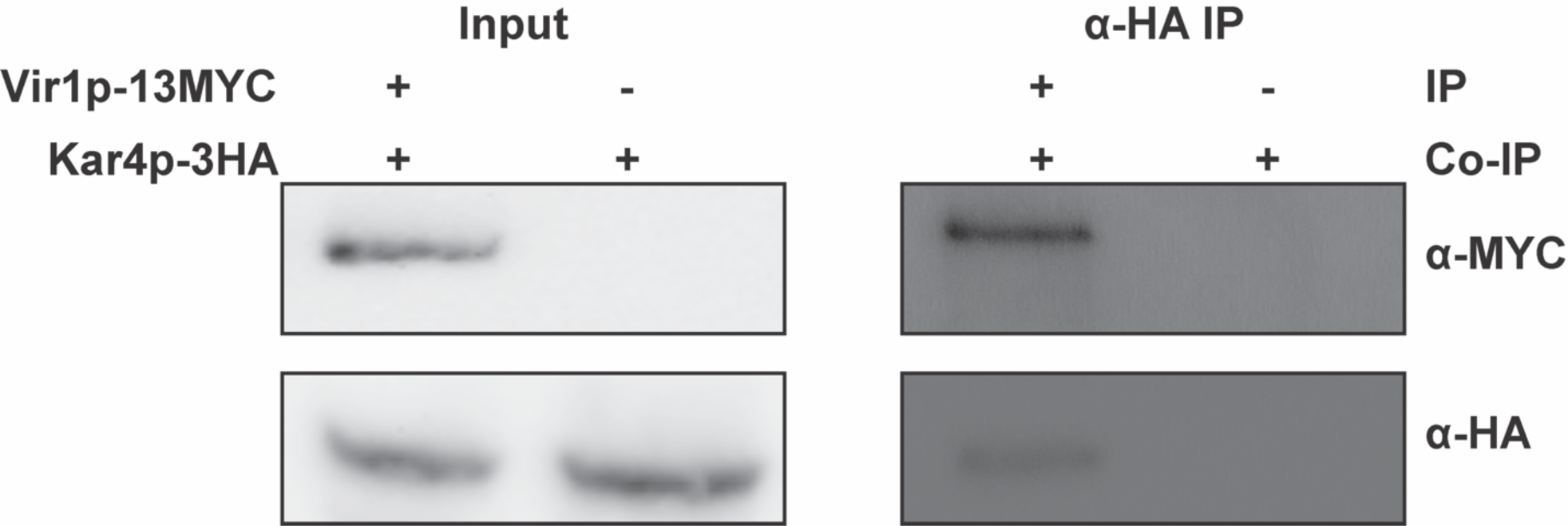
Vir1p and Kar4p interact. Further confirmation of the interaction between Vir1p and Kar4p. Co-IPs were conducted using both a tagged version of Vir1p (Vir1p-13MYC) and an untagged version in strains that both contained Kar4p-3HA. Interaction between the two proteins was only detected when both proteins were tagged.

**S Fig 2.**
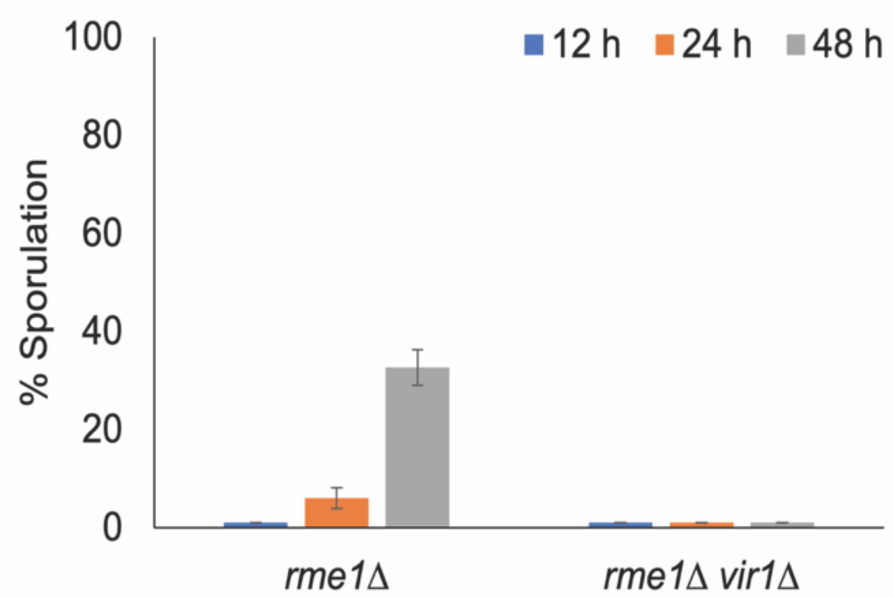
Loss of Rme1p does not permit sporulation in *vir1*Δ/Δ. Spores were counted at the indicated time points in *vir1*Δ*rme1*Δ/ *vir1*Δ*rme1*Δ and *rme1*Δ/ *rme1*Δ. All dyads, triads, and tetrads were counted. At least 100 cells were counted for each time point. Error bars represent the standard deviation of three biological replicates.

**S Fig 3.**
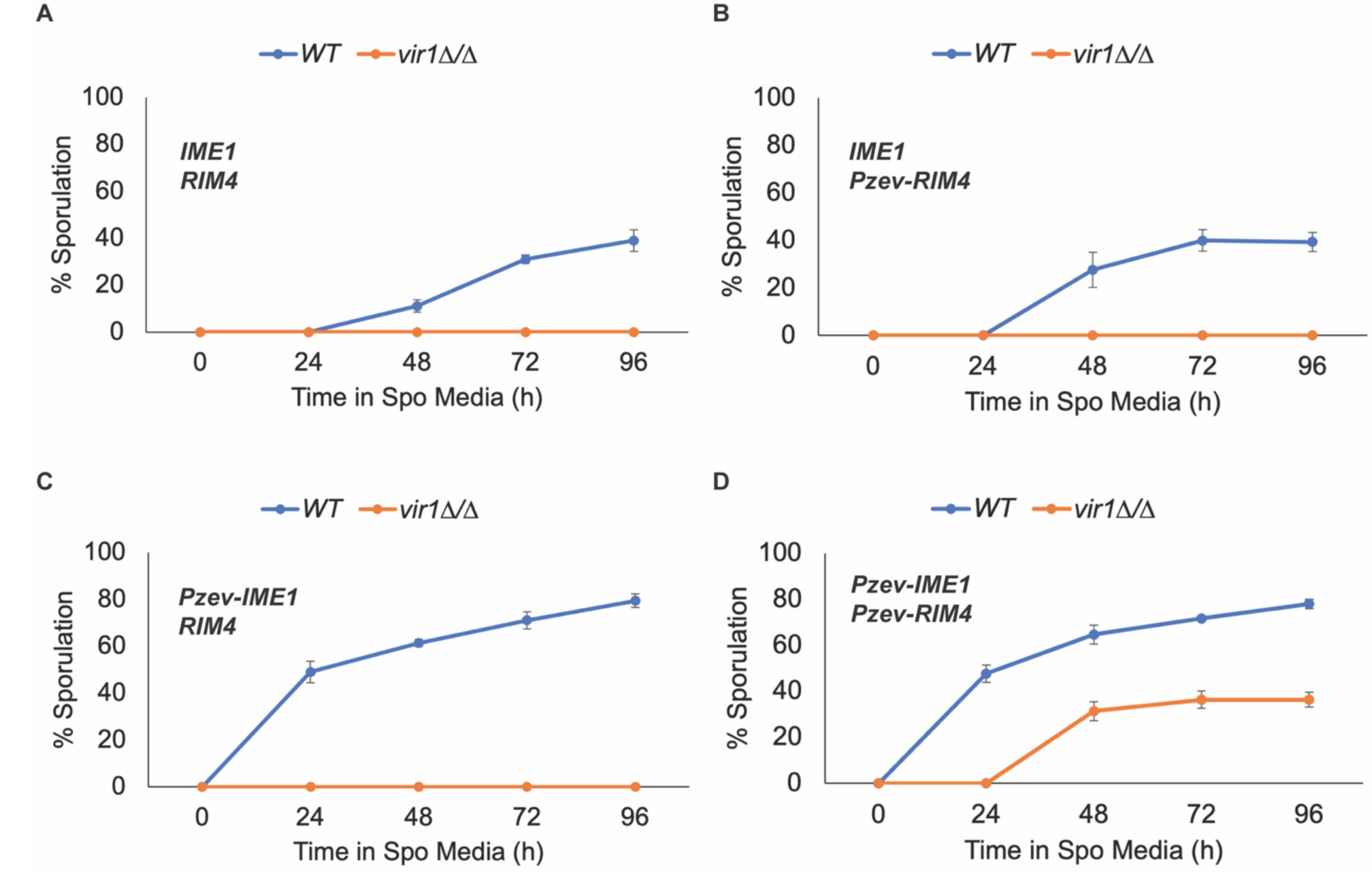
Sporulation of *vir1*Δ/Δ across an extended time course of meiosis. Percent of all tetrads, triads, and dyads summed across 96 hours in meiosis inducing conditions in wild-type and *vir1*Δ/Δ with either (A) *IME1/RIM4*, (B) *IME1/Pzev-RIM4*, (C) *Pzev-IME1/RIM4*, or (D) *Pzev-IME1/Pzev-RIM4*. Error bars are the standard deviation of three biological replicates.

**S Fig 4.**
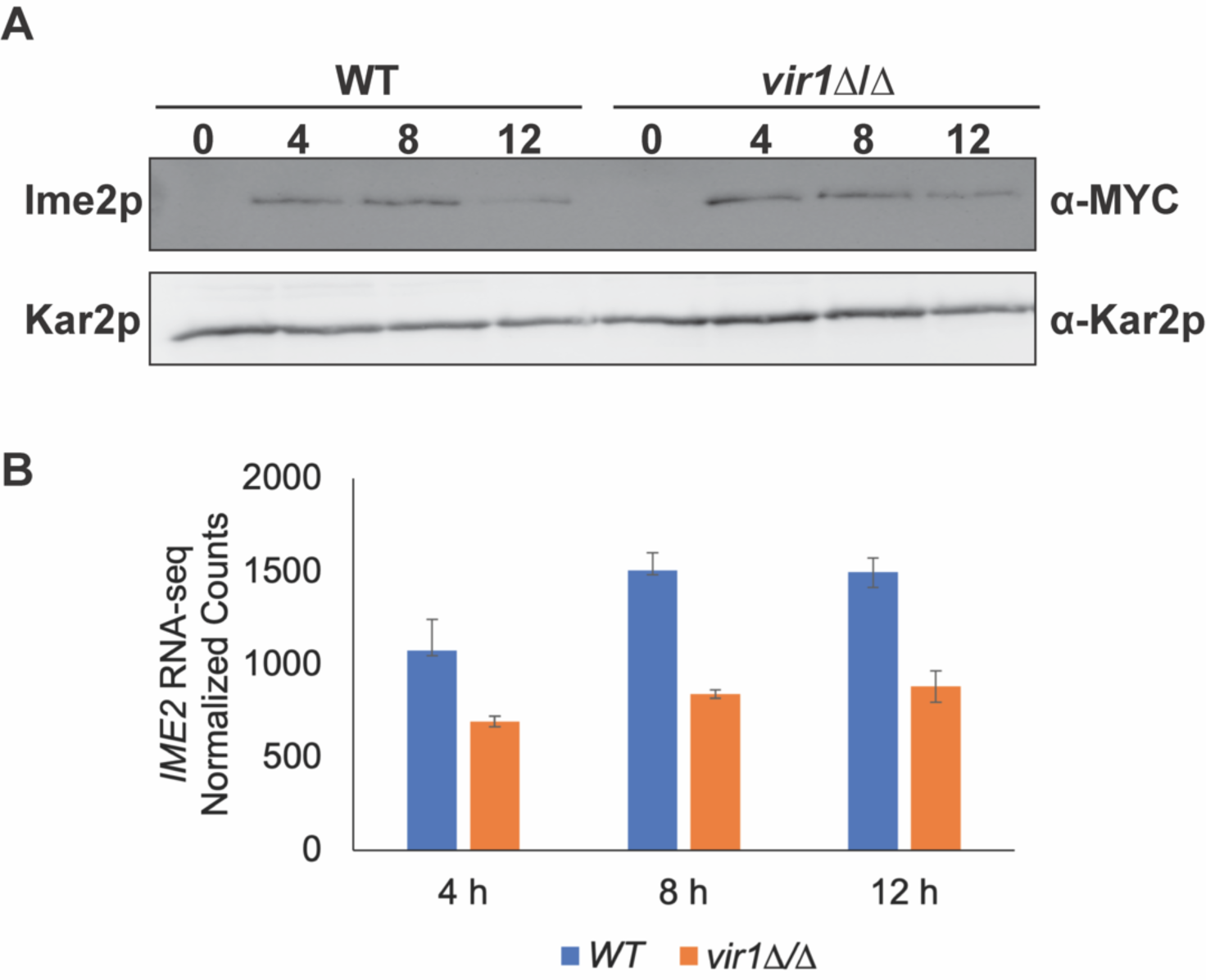
Loss of Vir1p does not impact Ime2p expression. (A) Western blots of Ime2p-13MYC across a meiotic time course in wild type and *vir1*Δ/Δ. Kar2p is used as a loading control. (B) *IME2* RNA-seq normalized counts from wild type and *kar4*Δ/Δ. Counts were normalized using the standard normalization method in DESeq2. Error bars represent standard deviation between two biological replicates.

**S Fig 5.**
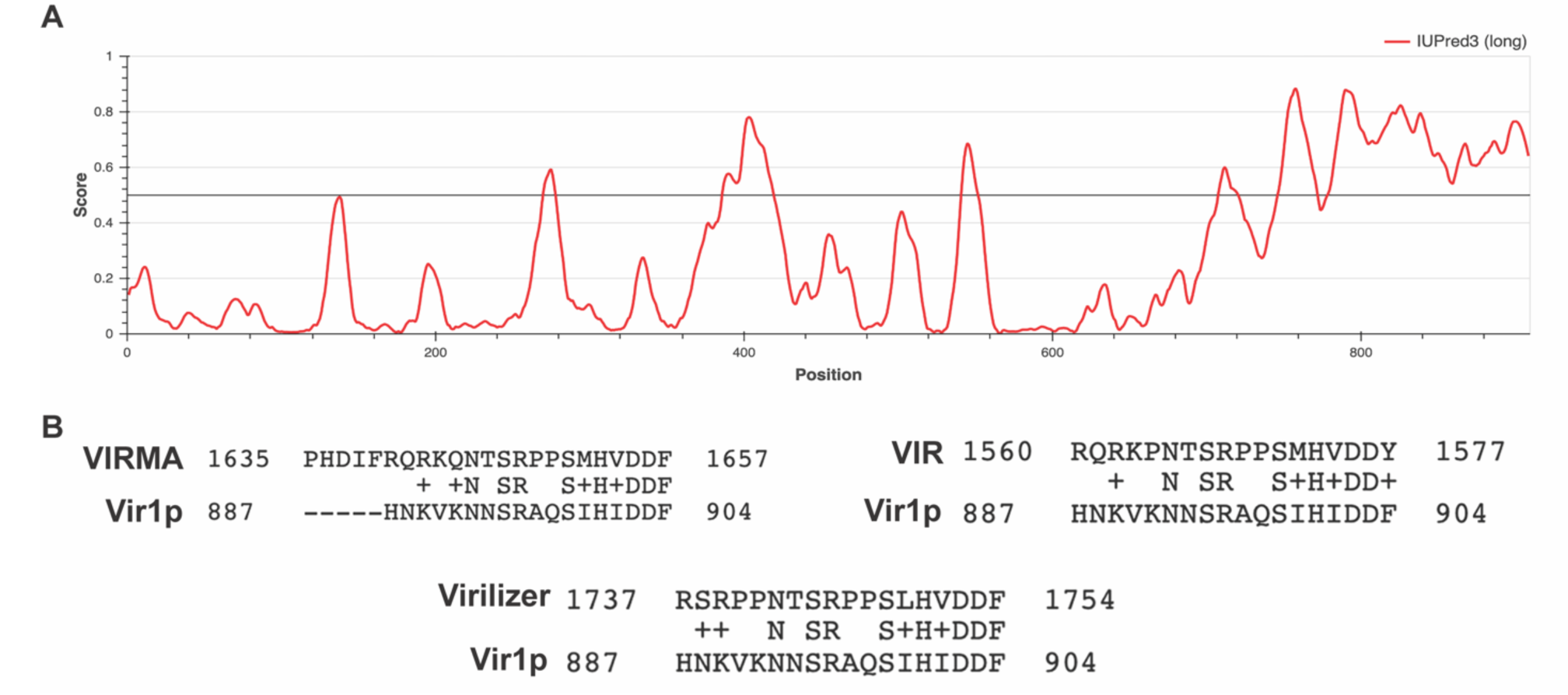
Vir1p’s C-terminus is predicted to be disordered. (A) Bokeh plot of IUPred3 predicted disordered regions of Vir1p. Standard settings were used in IUPred3. Regions above the solid black line have a higher probability of being disordered. (B) Blastp determined regions of conserved primary amino acid sequence between Vir1p and the VIRMA homologs.

**S Fig 6.**
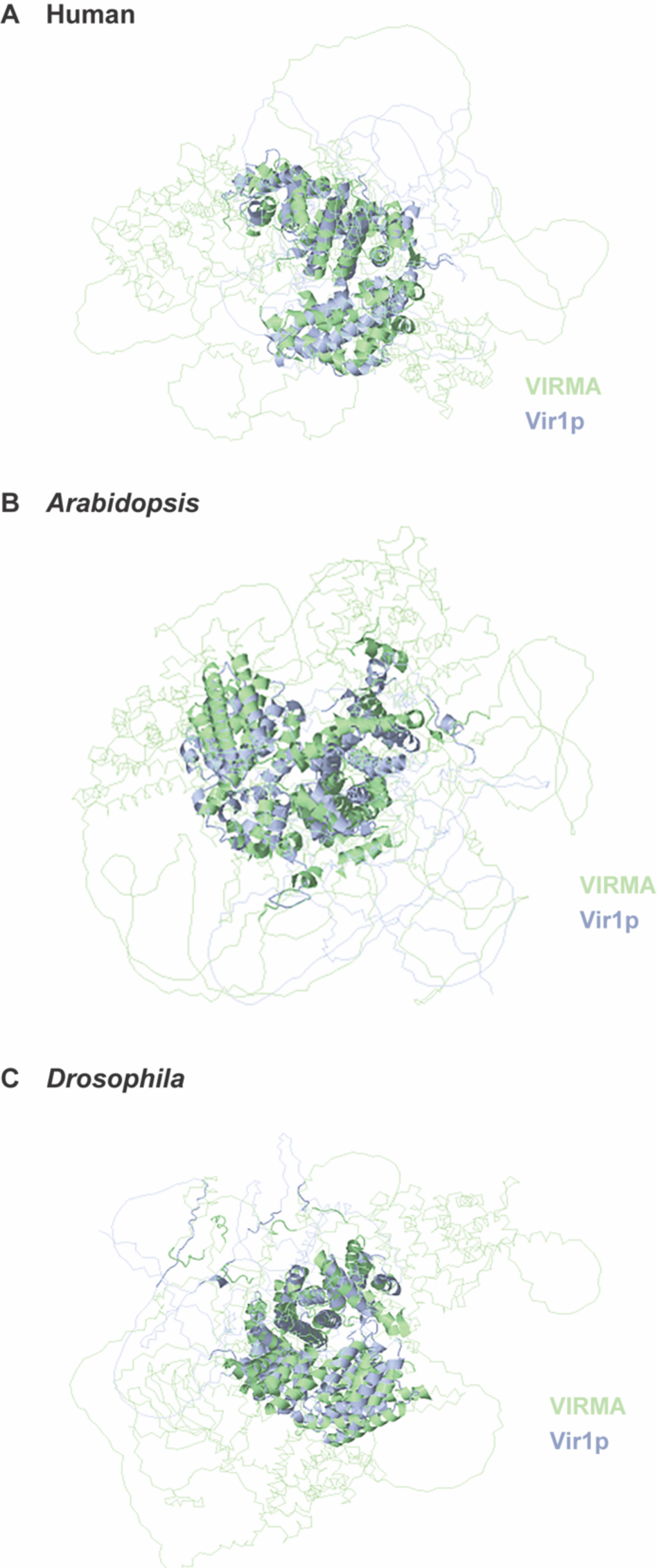
RaptorX comparison of Vir1p and VIRMA homologs. RaptorX predictions of conserved structure between Vir1p and VIRMA from (A) humans, (B) *Arabidopsis*, and (C) *Drosophila*. Highlighted regions of Vir1p (blue) and VIRMA (green) indicate regions of the two proteins that were predicted to have similar structure.

**S Fig 7.**
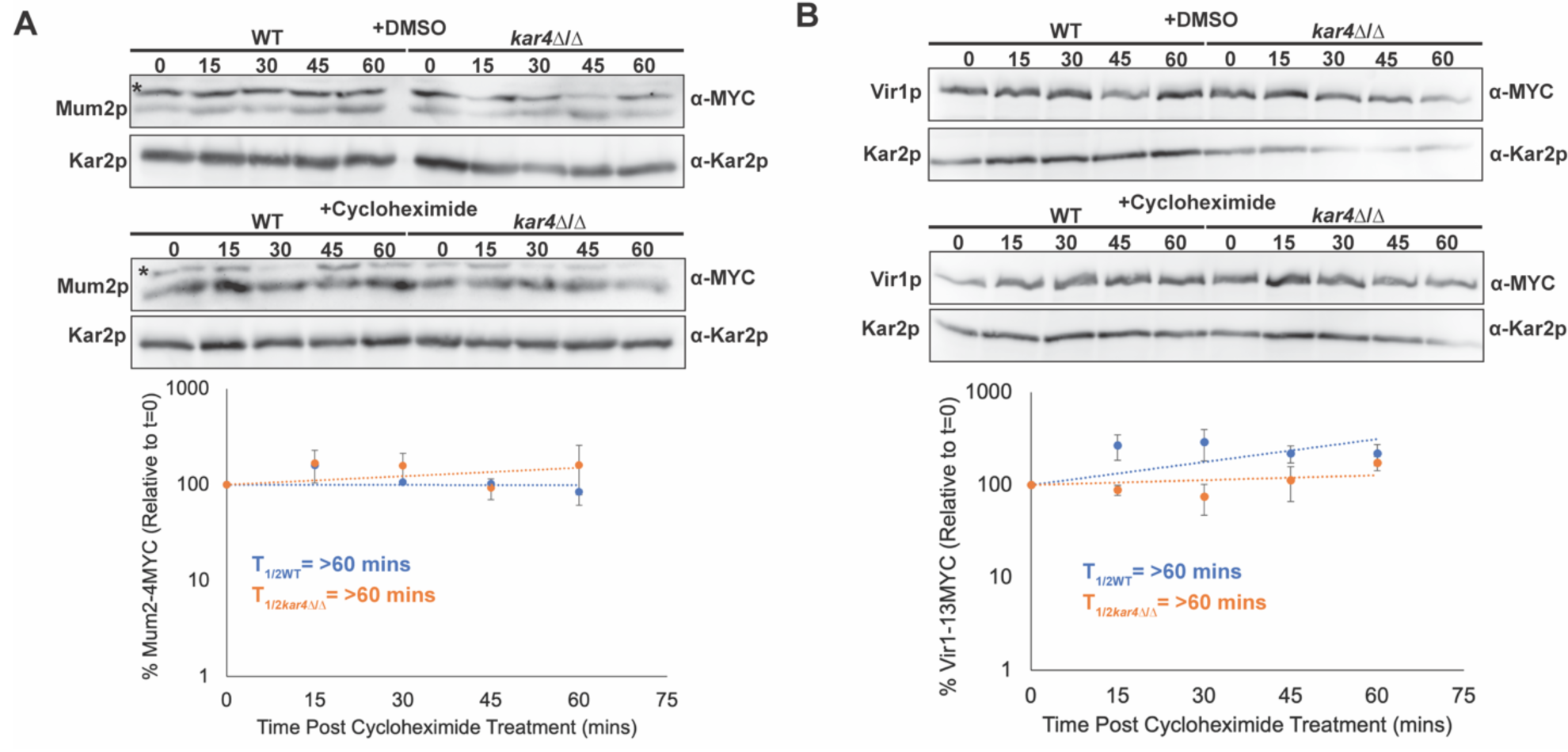
Loss of Kar4p does not impact the levels of other complex members. (A) Western blots of Mum2p-4MYC after four hours in meiosis inducing media with either 100 µM cycloheximide or an equivalent amount of DMSO in both wild type and *kar4*Δ/Δ. (Top) Mum2p-4MYC levels with DMSO. (Middle) Mum2p-4MYC levels with cycloheximide. Graph is the quantification of three biological replicates of the cycloheximide chase experiment. Kar2p is used as a loading control. “*” indicates a non-specific band. (B) Western blot of Vir1p-13MYC after four hours in meiosis inducing media with either 100 µM cycloheximide or an equivalent amount of DMSO in both wild type and *kar4*Δ/Δ. (Top) Vir1p-13MYC levels with DMSO. (Middle) Vir1p-13MYC levels with cycloheximide. Graph is the quantification of three biological replicates of the cycloheximide chase experiment. Kar2p is used as a loading control. Error bars in A and B are SEM of the three biological replicates.

**S Fig 8.**
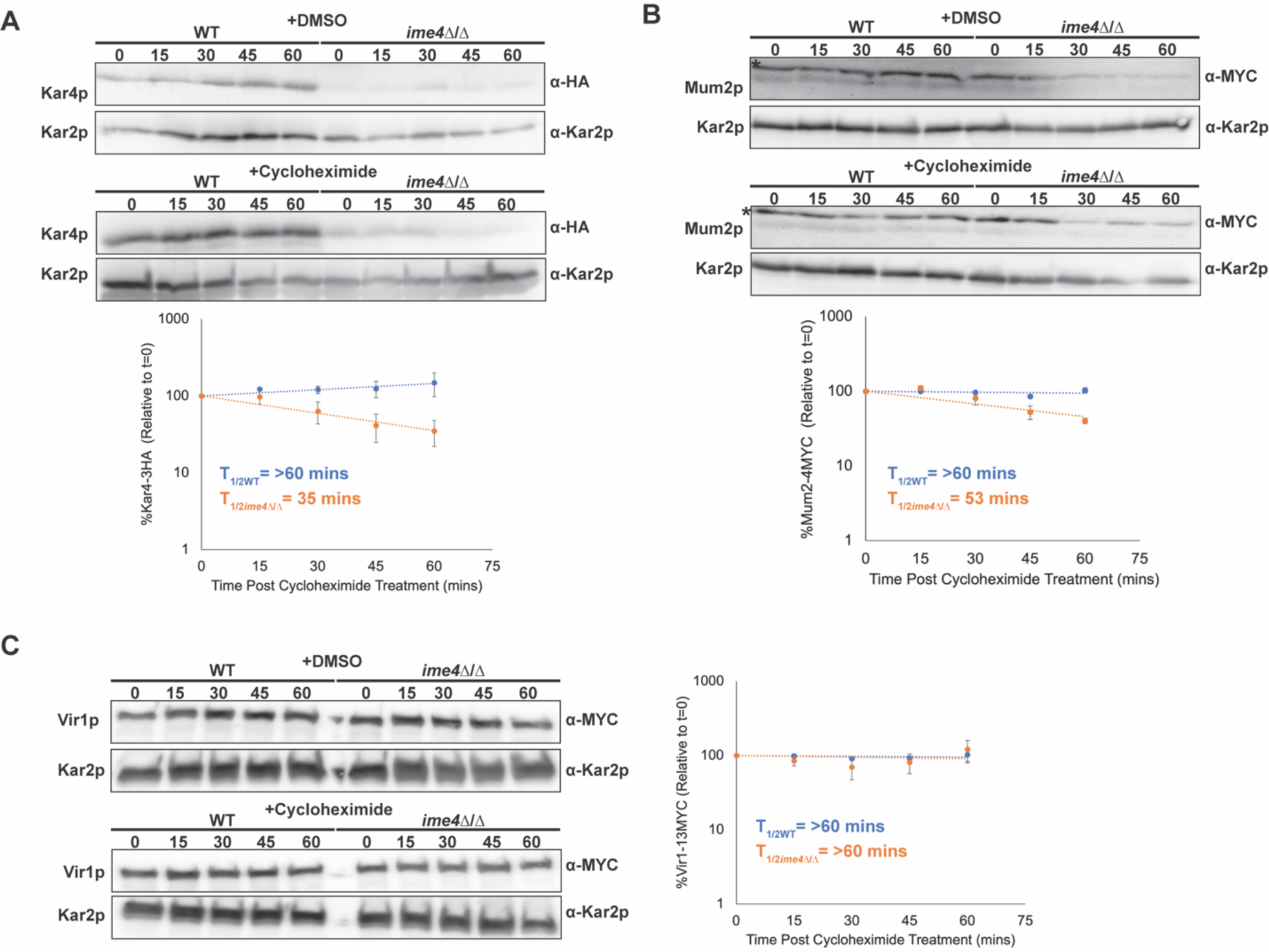
Loss of Ime4p results in increased turnover of Kar4p. (A) Western blots of Kar4p-3HA after four hours in meiosis inducing media with either 100 µM cycloheximide or an equivalent amount of DMSO in both wild type and *ime4*Δ/Δ. (Top) Kar4p-3HA levels with DMSO. (Bottom) Kar4p-3HA levels with cycloheximide. Graph is the quantification of three biological replicates of the cycloheximide chase experiment. Kar2p is used as a loading control. (B) Western blot of Mum2p-4MYC (bottom band) after four hours in meiosis inducing media with either 100 µM cycloheximide or an equivalent amount of DMSO in both wild type and *ime4*Δ/Δ. (Top) Mum2p-4MYC levels with DMSO. (Bottom) Mum2p-4MYC levels with cycloheximide. Graph is the quantification of three biological replicates of the cycloheximide chase experiment. Graph is the quantification of three biological replicates of the cycloheximide chase experiment. Kar2p is used as a loading control. “*” indicates a non-specific band. (C) Western blot of Vir1p-13MYC after four hours in meiosis inducing media with either 100 µM cycloheximide or an equivalent amount of DMSO in both wild type and *ime4*Δ/Δ. (Top) Vir1p-13MYC levels with DMSO. (Bottom) 3MYC-Ime4 levels with cycloheximide. Graph is the quantification of three biological replicates of the cycloheximide chase experiment. Kar2p is used as a loading control. Error bars in A, B, and C are SEM of the three biological replicates.

**S Fig 9.**
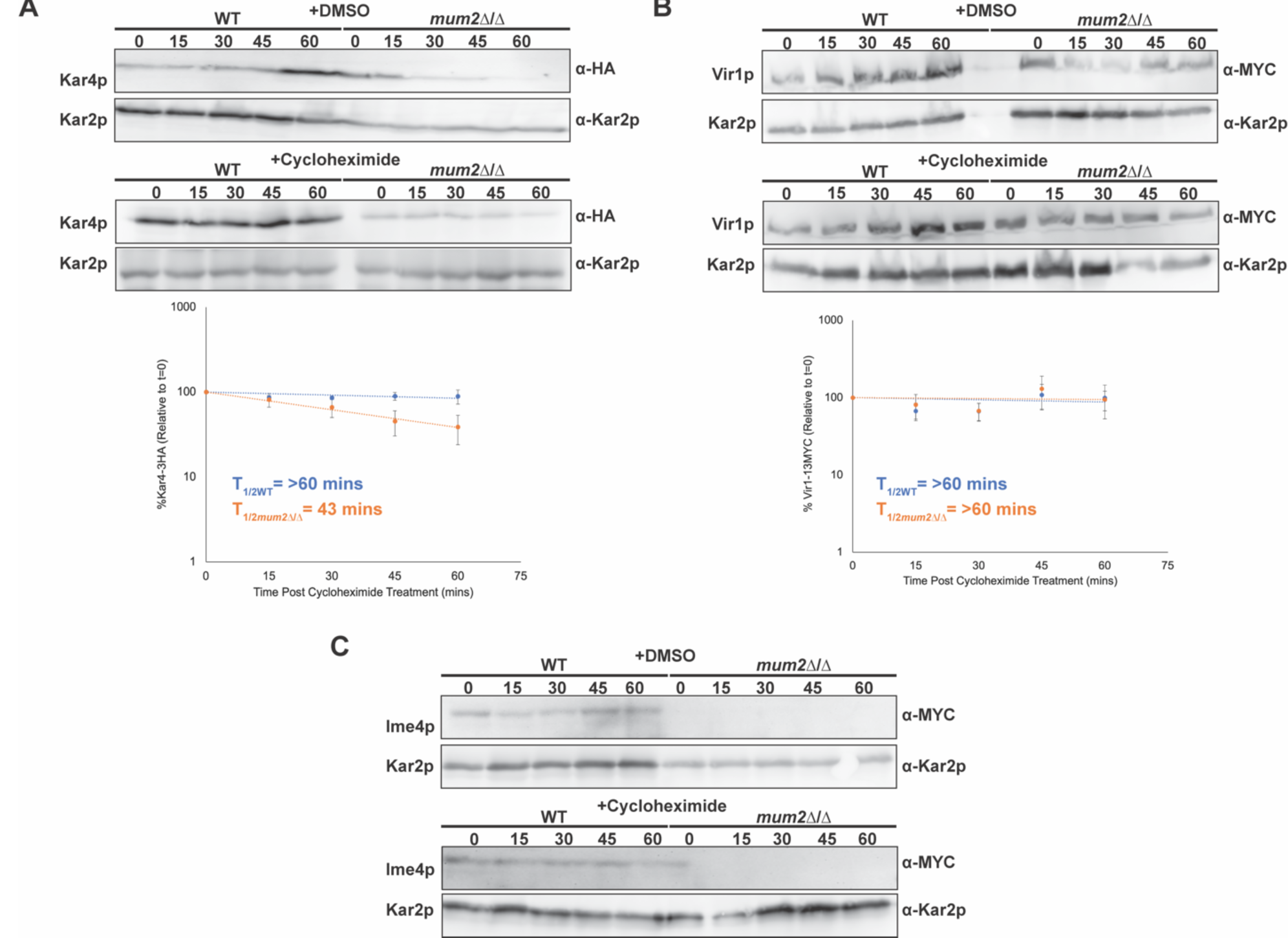
Loss of Mum2p results in reduced levels of all other complex members. (A) Western blots of Kar4p-3HA after four hours in meiosis inducing media with either 100 µM cycloheximide or an equivalent amount of DMSO in both wild type and *mum2*Δ/Δ. (Top) Kar4p-3HA levels with DMSO. (Bottom) Kar4p-3HA levels with cycloheximide. Graph is the quantification of three biological replicates of the cycloheximide chase experiment. Kar2p is used as a loading control. (B) Western blot of Vir1p-13MYC (bottom band) after four hours in meiosis inducing media with either 100 µM cycloheximide or an equivalent amount of DMSO in both wild type and *mum2*Δ/Δ. (Top) Vir1p-13MYC levels with DMSO. (Bottom) Vir1p-13MYC levels with cycloheximide. Graph is the quantification of three biological replicates of the cycloheximide chase experiment. Kar2p is used as a loading control. “*” indicates a non-specific band. Error bars in A and B are SEM of the three biological replicates. (C) Western blot of 3MYC-Ime4p after four hours in meiosis inducing media with either 100 µM cycloheximide or an equivalent amount of DMSO in both wild type and *mum2*Δ/Δ. (Top) 3MYC-Ime4p levels with DMSO. (Bottom) 3MYC-Ime4p levels with cycloheximide. Kar2p is used as a loading control.

**Table S1.** Strains used for this study. Auxotrophic markers are standard BY strain alleles.

| Strain Name | Genotype | Strain | Reference |
| --- | --- | --- | --- |
| MY 16019 | *ura3/" leu2/" his3/" lys2/+ met15/+ kar4::KANMX/" KFC1-13MYC::HIS3/+ [KAR4-3HA URA3]* | S288c |  |
| MY 16232 | *ura3/" leu2/" his3/" lys2/+ met15/+ MUM2-3HA::HIS3/+ kfc1::KANMX/KFC1-13MYC::HIS3* | S288c |  |
| MY 16236 | *ura3/" leu2/" his3/" lys2/+ met15/+ MUM2-3HA::HIS3/+ kfc1::KANMX/+* | S288c |  |
| MY 16327 | *ura3/" leu2/" his3/" lys2/+ met15/+ KFC1-13MYC-HIS3/kfc1::KANMX IME4-3HA::HIS3/+ kar4::KANMX/+* | S288c |  |
| MY 16328 | *ura3/" leu2/" his3/" lys2/+ met15/+ IME4-3HA::HIS3/+ kfc1::KANMX/+* | S288c |  |
| MY 16408 | *ura3/" leu2/" his3/" lys2/+ met15/+ SLZ1-3HA::HIS3/+ KFC1-13MYC::HIS3/+ kfc1::KANMX/+ kar4::KANMX/+* | S288c |  |
| MY 16409 | *ura3/" leu2/" his3/" lys2/+ met15/+ SLZ1-3HA::HIS3/+ kfc1::KANMX/+* | S288c |  |
| MY 16325 | *ura3/" lys2/" ho::LYS2/" arg4-BglIII/ARG4 leu2::HisG/LEU2* | SK1 |  |
| MY 16326 | *ura3/" lys2/" ho::LYS2/" arg4-BglIII/ARG4 leu2::HisG/LEU2 ime4::URA3/"* | SK1 |  |
| MY 16369 | *ho::LYS2/" arg4-BglIII/+ +/leu2::hisG lys2-SK1/" ura3::hisG/" kfc1::URA3/"* | SK1 |  |
| MY 16563 | *leu2/" his3/" ura3/" met15/+ lys2/+ 3xFLAG-RME1/"* | S288c |  |
| MY 16575 | *leu2/" his3/" ura3/" met15/+ lys2/+ 3xFLAG-RME1/" kfc1::HPHMX/"* | S288c |  |
| MY 16550 | *his3/" leu2/" ura3/" lys2/+, met15/+ GFP-IME1/"* | S288c |  |
| MY 16574 | *his3/" leu2/" ura3/" lys2/+ met15/+ GFP-IME1/GFP-IME1 kfc1::HPHMX/"* | S288c |  |
| MY 16294 | *URA3::GFP-TUB1/+ SPC42-mCherry::HIS3/+ kar4::URA3/+ HPHMX::Pz3ev-IME1/+ leu2::ACT1PR-3ZEV-NATMX/leu2 his3/+ ura3/" met15/+* | S288c |  |
| MY 16365 | *URA3::GFP-TUB1/+ SPC42-mCherry::HIS3/+ kfc1::KANMX/kfc1::URA3 HPHMX::Pz3ev-IME1/+ leu2::ACT1PR-3ZEV-NATMX/leu2 his3/+ ura3/" met15/+* | S288c |  |
| MY 16616 | *his3/" leu2/leu2::pACT1-Z3EV-NATMX +/lys2 met17/+ ura3/"* | S288c |  |
| MY 16618 | *his3/" leu2/leu2::pACT1-Z3EV-NATMX +/lys2 met17/+ ura3/" kfc1::KANMX/"* | S288c |  |
| MY 16609 | *his3/" leu2/" ura3/" lys2/+ met15/+ rme1::KANMX/" kfc1::HPHMX/"* | S288c |  |
| MY 16456 | *rme1::KANMX/" leu2/" his3/" ura3/" lys2/+ met15/+* | S288c |  |
| MY 16532 | *HPHMX::PZ3EV-RIM4/+ leu2::act1pr-Z3EV-NATMX/leu2 ura3/" his3/+ lys2/+* | S288c |  |
| MY 16533 | *HPHMX::PZ3EV-IME1/+ HPHMX::PZ3EV-Rim4/+ leu2::act1pr-Z3EV-NATMX/" ura3/"* | S288c |  |
| MY 16534 | *HPHMX::PZ3EV-IME1/+ leu2::act1pr-Z3EV-NATMX/leu2 ura3/" his3/+ lys2/+* | S288c |  |
| MY 16293 | *HPHMX::PZ3EV-IME1/+ leu2::act1pr-Z3EV-NATMX/leu2 ura3/" his3/+ lys2/+ kfc1::KANMX/kfc1::URA3* | S288c |  |
| MY 16073 | *HPHMX::PZ3EV-IME1/+ HPHMX::PZ3EV-Rim4/+ leu2::act1pr-Z3EV-NATMX/" ura3/" kfc1::URA3/"* | S288c |  |
| MY 16208 | *HPHMX::PZ3EV-IME1/+ leu2::act1pr-Z3EV-NATMX/leu2, ura3/", his3/+, met15/+ kar4::kanMX4/+ GAS4-3HA::HIS3/+* | S288c |  |
| MY 16209 | *HPHMX::PZ3EV-IME1/+ leu2::act1pr-Z3EV-NATMX/leu2, ura3/", his3/+, met15/+ kar4::kanMX4/+ SPS2-3HA::HIS3/+* | S288c |  |
| MY 16077 | *HPHMX::PZ3EV-RIM4/+ leu2::act1pr-Z3EV-NATMX/leu2 ura3/" his3/+ lys2/+ kfc1::URA3/kfc1::KANMX* | S288c |  |
| MY 16215 | *SPS2-3HA::HIS3/+ HPHMX::PZ3EV-IME1/+ leu2::act1pr-Z3EV-NATMX/leu2 ura3/" his3/+ met15/+ kfc1::KANMX/kfc1::URA3* | S288c |  |
| MY 16219 | *GAS4-3HA::HIS3/+ HPHMX::PZ3EV-IME1/+ leu2::act1pr-Z3EV-NATMX/leu2 ura3/" his3/+ met15/+ kfc1::KANMX/kfc1::URA3* | S288c |  |
| MY 16292 | *SPS2-3HA::HIS3/+ kar4::KANMX/+ kfc1::URA3/kfc1::LEU2 HPHMX::PZ3EV-IME1/+ HPHMX::PZ3EV-Rim4/+ leu2::act1pr-Z3EV-NATMX/" met15/+* | S288c |  |
| MY 16371 | *GAS4-3HA::HIS3/+ kar4::KANMX/+ kfc1::URA3/kfc1::LEU2 HPHMX::PZ3EV-IME1/+ HPHMX::PZ3EV-Rim4/+ leu2::act1pr-Z3EV-NATMX/" met15/+* | S288c |  |
| MY 16572 | *his3/" leu2/" ura3/" lys2/+ met15/+ IME2-13MYC::KANMX/+* | S288c |  |
| MY 16603 | *his3/" leu2/" ura3/" lys2/+ met15/+ IME2-13MYC::KANMX/+ kfc1::HPHMX/"* | S288c |  |
| MY 16611 | *kfc1::HPHMX/kfc1::URA3 his3/+ lys2/+ ura3/" IME2-13MYC::KANMX/+ HPHMX::PZ3EV-IME1/+ leu2::act1pr-Z3EV-NATMX/leu2* | S288c |  |
| MY 16628 | *kfc1::URA3/" IME2-13MYC::KANMX/+ HPHMX::PZ3EV-IME1/+ HPHMX::PZ3EV-RIM4/+ leu2::act1pr-Z3EV-NATMX/" ura3/+ his3/+ met15/+* | S288c |  |
| MY 16508 | *leu2/" his3/" ura3/" met15/+ lys2/+ MUM2-4MYC::HIS3/" kfc1::KANMX/"* | S288c |  |
| MY 16522 | *leu2/" his3/" ura3/" met15/+ lys2/+ MUM2-4MYC::HIS3/"* | S288c |  |
| MY 16518 | *his3/" leu2/" ura3/" met15/+ lys2/+ [KAR4-3HA URA3]* | S288c |  |
| MY 16519 | *kfc1::KANMX/" his3/" leu2/" ura3/" met15/+ lys2/+ [KAR4-3HA URA3]* | S288c |  |
| MY 16542 | *leu2/" his3/" ura3/" met15/+ lys2/+ ime4::LEU2/" [KAR4-3HA URA3]* | S288c |  |
| MY 16374 | *kar4::KANMX/+ leu2/" his3/" ura3/" lys2/+ met15/+ KFC1-13MYC::HIS3/+ mum2::KANMX/mum2::LEU2 [KAR4-3HA URA3]* | S288c |  |
| MY 16256 | *leu2/" his3/" ura3/" met15/+ lys2/+ kar4::KanMX/" MUM2-4MYC::HIS3/+ [URA3]* | S288c |  |
| MY 16257 | *leu2/" his3/" ura3/" met15/+ lys2/+ kar4::KanMX/" MUM2-4MYC::HIS3/+ [KAR4-3HA URA3]* | S288c |  |
| MY 16625 | *ura3/" leu2/" his3/" lys2/+ met15/+ ime4::NATMX/" MUM2-4MYC::HIS3/"* | S288c |  |
| MY 16626 | *ura3/" leu2/" his3/" lys2/+ met15/+ kar4::KANMX/+ KFC1-13MYC::HIS3/+* | S288c |  |
| MY 16627 | *ura3/" leu2/" his3/" lys2/+ met15/+ kar4::KANMX/" KFC1-13MYC::HIS3/+* | S288c |  |
| MY 16437 | *kar4::KANMX/+ leu2/" his3/" ura3/" lys2/+ met15/+ VIR1-13MYC::HIS3/+* | S288c |  |
| MY 16438 | *kar4::KANMX/+ leu2/" his3/" ura3/" lys2/+ met15/+ ime4::NATMX/ime4::LEU2 VIR1-13MYC::HIS3/+* | S288c |  |
| MY 16613 | *3MYC-IME4/" mum2::KANMX/" ho::LYS2/" lys2-SK1/"* | SK1 |  |
| SAy914 | *3MYC-IME4/" ho::LYS2/" lys2-SK1/"* | SK1 | Agarwala et al.<br>2012 |
| MY 16629 | *ura3/" lys2-SK1/" ho::LYS2/" arg4-BgIII/+ leu2::HisG/+ 3MYC-IME4/"* | SK1 |  |
| MY 16630 | *ho::LYS2/" arg4-BgIII/+ +/leu2::hisG lys2-SK1/" ura3::hisG/" kfc1::URA3/" 3MYC-IME4/"* | SK1 |  |

**Table S2.** Plasmids used in this paper.

| Strain | Markers | Source |
| --- | --- | --- |
| pMR<br>2654 | *CEN ARS URA3 KAR4-3HA* |  |
| pAFS<br>125 | *GFP-TUB1 URA3 Amp* | Straight et al.<br>1997 |
| pRB<br>3483 | *hphMx-Pz3ev CEN Amp* | McIsaac et al.<br>2014 |

